# Coherent feedback leads to robust background compensation in oscillatory and non-oscillatory homeostats

**DOI:** 10.1101/2023.05.30.542992

**Authors:** Melissa Nygård, Peter Ruoff

## Abstract

When in an integral feedback controller a step perturbation is applied at a constant background, the controlled variable (described here as *A*) will in general respond with decreased response amplitudes Δ*A* as backgrounds increase. The controller variable *E* will at the same time provide the necessary compensatory flux to move *A* back to its set-point. A typical example of decreased response amplitudes at increased backgrounds is found in retinal light adaptation. Due to remarks in the literature that retinal light adaptation would also involve a compensation of backgrounds we became interested in conditions how background compensation could occur. In this paper we describe how background influences can be robustly eliminated. When such a background compensation is active, oscillatory controllers will respond to a defined perturbation with always the same (damped or undamped) frequency profile, or in the non-oscillatory case, with the same response amplitude Δ*A*, irrespective of the background level. To achieve background compensation we found that two conditions need to apply: (i) an additional set of integral controllers (here described as *I*_1_ and *I*_2_) have to be employed to keep the manipulated variable *E* at a defined set-point, and (ii), *I*_1_ and *I*_2_ need to feed back to the *A*-*E* signaling axis directly through the controlled variable *A*. In analogy to a similar feedback applied in quantum control theory, we term these feedback conditions as ‘coherent feedback’. When analyzing retinal light adaptations in more detail, we find no evidence in the presence of background compensation mechanisms. Although robust background compensation, as described theoretically here, appears to be an interesting regulatory property, relevant biological or biochemical examples still need to be identified.

## Introduction

Homeostatic mechanisms play important roles in physiology and in the adaptation of organisms to their environments [1]. For example in the vertebrate retina, photoreceptor cells contain negative feedback loops which participate in light adaptation [2–5]. A hallmark of vertebrate photoadaptation is that resetting kinetics accelerate and response amplitudes decrease as backgrounds increase [5, 6]. This behavior is seen in Fig 1 for a macaque monkey’s rod cell response towards a single light flash applied at different background light levels.

**Fig 1.**
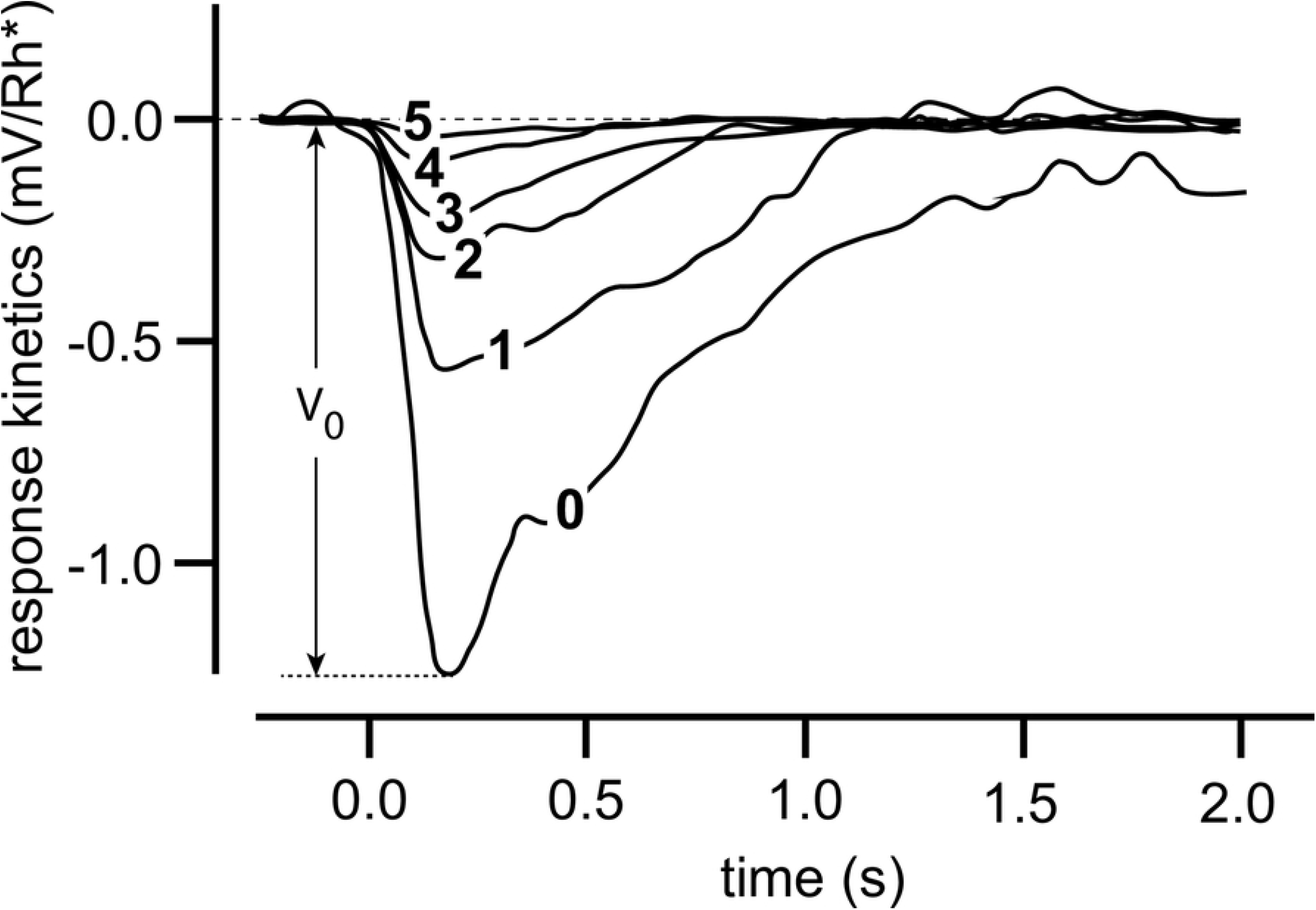
Light adaptation in a Macaque monkey’s rod cell. 10 ms light flashes were applied to different light background intensities. Background intensities (in photons *μ*m^*−*2^s^*−*1^) were: **0**, 0; **1**, 3.1; **2**, 12; **3**, 41; **4**, 84; **5**, 162. The influence of the background on the response amplitude and the speed of resetting is clearly seen. *V*_0_ is the response amplitude for background **0**. Redrawn and modified after Fig 2A from Ref [7]. For a theoretical description of this behavior see Ref [5] and references therein.

Another retinal light adaptation example is shown in Fig 2. Here, the mean maximum firing rates of a cat ganglion cell was measured with respect to different step light perturbations (test spot luminance) which are applied at six different backgrounds [8].

**Fig 2.**
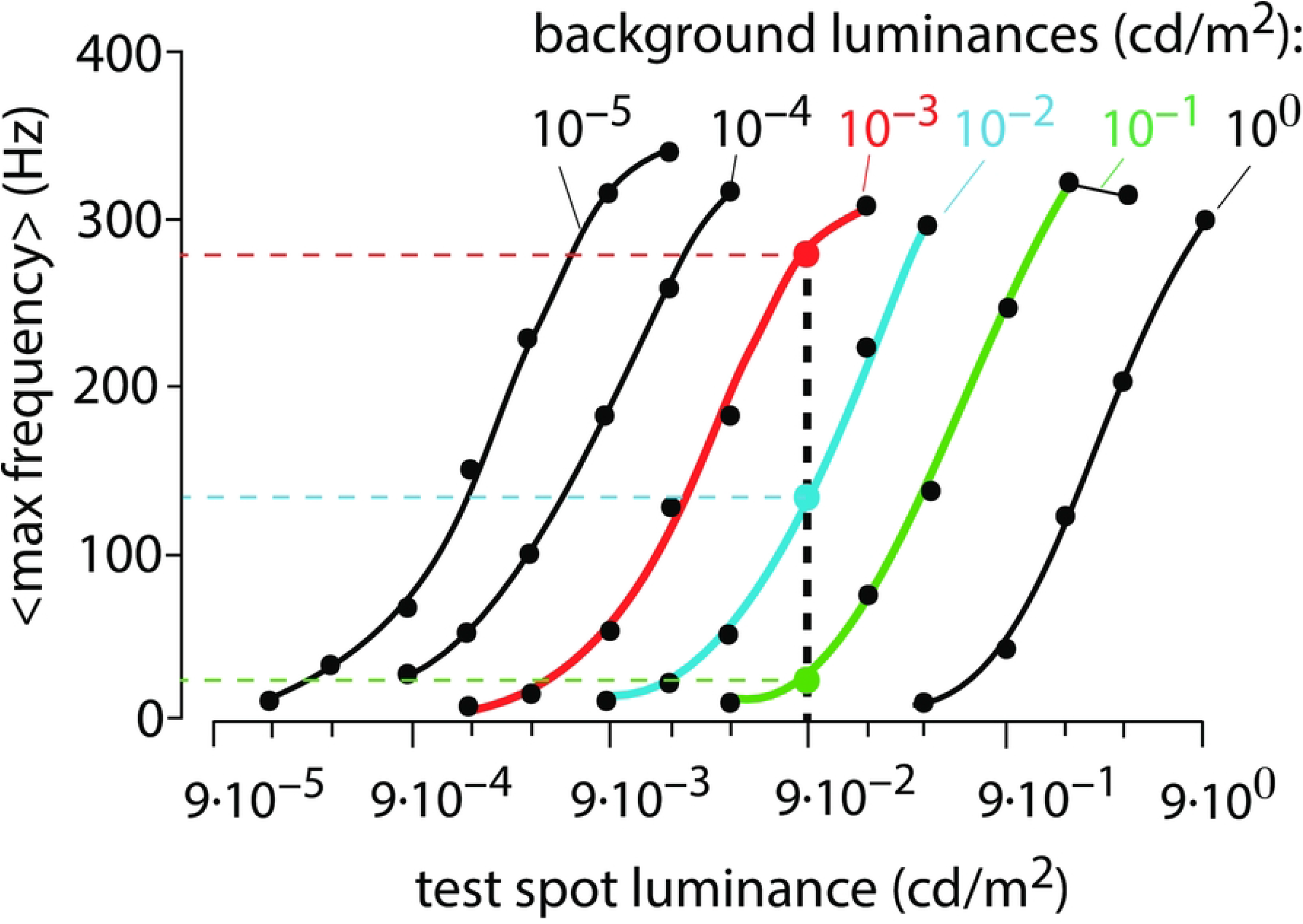
Light adaptation of an on-center ganglion cell in the cat retina (redrawn from Fig 8, Ref [8]). Averaged maximum ganglion cell frequencies are shown as a function of six different background illuminations in response to applied light step perturbations (test spot luminance). The three colored curves show the averaged maximum frequencies at background illuminations 10^*−*3^, 10^*−*2^ and 10^*−*1^ cd/m^2^. A test spot luminance (perturbation) of 9*×*10^*−*2^ cd/m^2^ is indicated as the vertical dashed black line. The colored intersection points and vertical dashed lines indicate that for this perturbation strength the maximum mean response frequency decreases with increasing background illumination.

In Kandel et al. [2] it was commented (see page 540, section *Light Adaptation Is Apparent in Retinal Processing and Visual Perception*) that Fig 2 would indicate a compensation of the background illumination and thereby causing the same response due to a lateral shifting along the perturbation (test spot luminance) axis. Based on this comment we became interested in mechanisms which would allow to compensate for background levels and thereby give the same response for a given perturbation irrespective of the applied background.

In this paper we present results on how such a robust background compensation can be achieved in both oscillatory and non-oscillatory homeostats.

The paper is structured in the following way: We first show that a feedback type similar to what quantum physicists have termed ‘coherent feedback’ [9, 10] is required to obtain background compensation in both oscillatory and non-oscillatory homeostats. For oscillatory homeostats we show that coherent feedback control leads to, besides background compensation, also to frequency control. In fact, robust frequency control was previously observed by us [11], but without having recognized the background-compensating property of coherent feedback. For non-oscillatory controllers or homeostats with damped oscillations, coherent feedback leads to conserved response profiles in the controlled variables, independent of an applied background. We then look at the situation of a ‘incoherent feedback’, where background compensation is lost, but oscillatory homeostats may still show robust frequency control. Finally we analyze photoreceptor responses in terms of a Michaelis-Menten model [3] and show that parallel lines as in Fig 2 or as log-log plots do not require the postulation of background compensation or additional adaptation mechanisms.

## Materials and methods

Computations were performed with the Fortran subroutine LSODE [12], which can be downloaded from https://computing.llnl.gov/projects/odepack. Graphical output was generated with gnuplot (www.gnuplot.info) and annotated with Adobe Illustrator (https://www.adobe.com/).

To make notations simpler, concentrations of compounds are denoted by compound names without square brackets. Time derivatives are generally indicated by the ‘dot’ notation. Rate parameters are in arbitrary units (au) and are presented as *k*_*i*_’s (*i*=1, 2, 3, …) irrespective of their kinetic nature, i.e. whether they represent turnover numbers, Michaelis constants, or inhibition/activation constants. To allow readers to redo calculations, the supporting information S1 Programs contains python scripts for a set of selected results.

### Usage of integral control

In the calculations robust homeostasis of concentrations and frequencies is achieved by implementing integral control into the negative feedbacks, a concept which comes from control engineering [13–16], and has been indicated to occur in and being applied to biological systems [17–21]. Briefly, in integral control the difference (also termed error) between the actual concentration of a controlled variable *A* and its set-point is integrated in time. The integrated error can then be used to compensate precisely for step-wise perturbations [15, 16]. Fig 3a shows the control scheme of integral control. An example is given in panel b using ‘motif 2’, one of eight basic feedback loops [22]. Panel c shows how zero-order removal of controller species *E* leads to integral control with a defined set-point of the controlled variable *A*.

**Fig 3.**
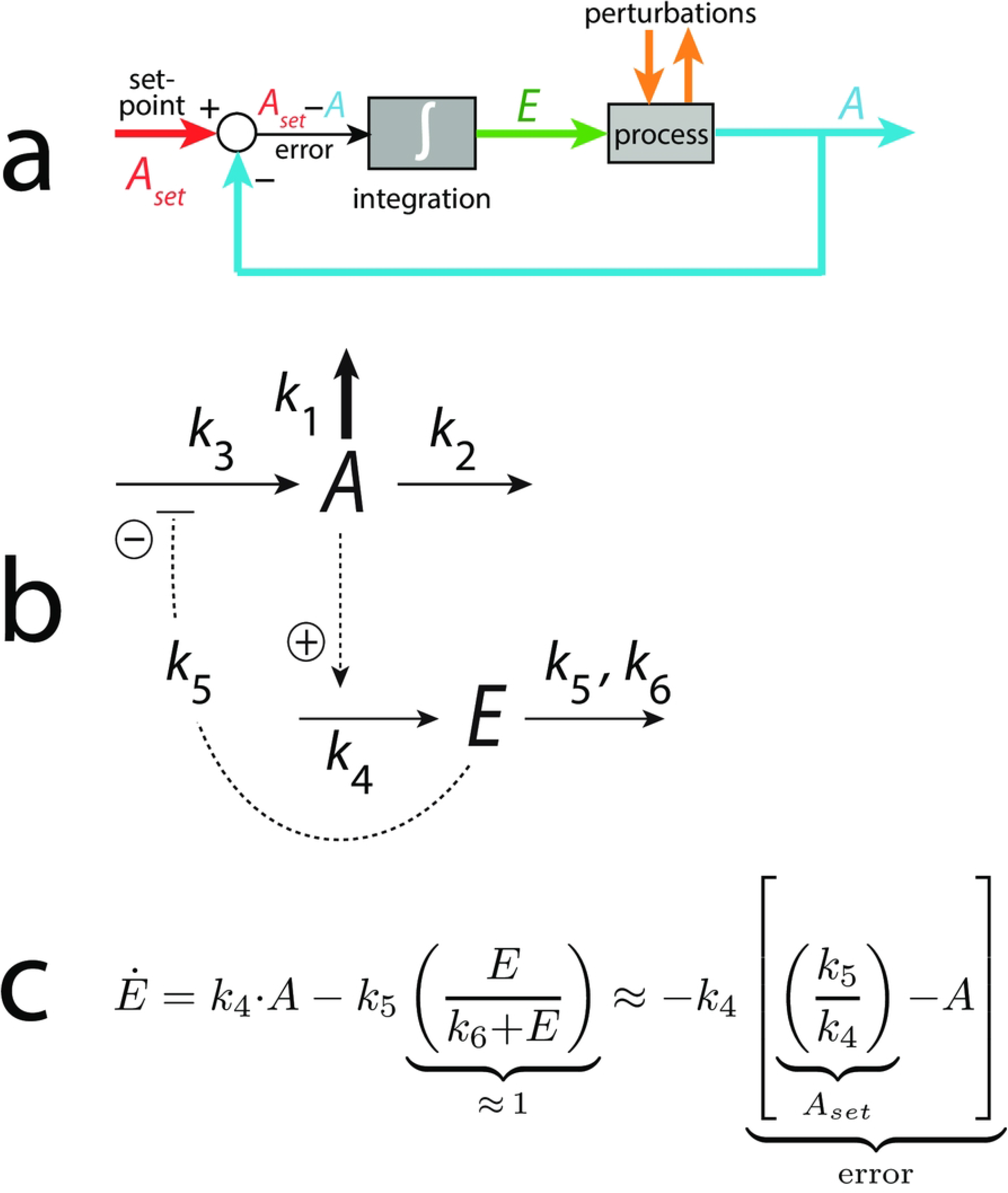
Principle of integral control. Panel a: The controlled variable *A* (outlined in blue) is compared with its set-point and the difference/error (*A*_*set*_*−A*) is integrated. This leads to the integrated error *E*, which is able to compensate precisely for step-wise perturbations [15]. Panel b: Basic negative feedback loop (motif 2, [22]). Solid lines are chemical reactions, while dashed lines represent activations (plus sign) and inhibitions (negative sign). Panel c: Rate equation of controller *E*. The zero-order removal of *E* introduces integral control. The set-point for *A* is given as *k*_5_/*k*_4_, and the concentration of *E* becomes proportional to the integrated error. For details, see for example Ref [22].

In the below calculations we have used zero-order kinetics to implement integral control [17, 22–24]. However, it should be mentioned that there are other kinetics conditions to achieve integral control, such as antithetic control [21, 25, 26], which generally will show identical resetting behaviors as in zero-order control [5, 26]. Also, autocatalytic reactions can be used to obtain integral control [27–29], which will generally show much faster resetting kinetics in comparison when integral control is introduced by zero-order kinetics [30, 31].

## Results and discussion

### Background compensation in negative feedback oscillators by coherent feedback

In this section we describe feedback conditions which can achieve background compensation in oscillatory homeostats. The type of oscillators we here focus on show frequency homeostasis due to a two-layered negative feedback structure. The center negative feedback layer ensures that the time average value of a controlled variable *A*, defined by Eq 1, is kept robustly at a certain set-point by a controller species *E* via integral control.

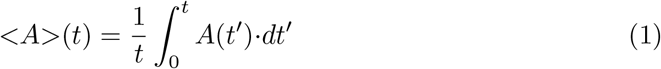

A second ‘outer’ negative feedback layer keeps on its side the time average value of *E*, i.e. *<E>* (Eq 2), under robust homeostatic control by two additional controller variables *I*_1_ and *I*_2_.

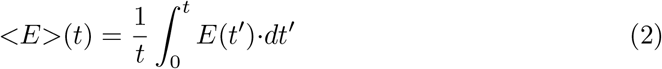

We previously showed [11] that these two-layered negative feedback structures enable robust frequency homeostasis. Here we now report the additional finding that when the *I*_1_ and *I*_2_ controllers feed back directly via *A* to control *E*, the oscillator has the capability to neutralize backgrounds. In analogy to a closely related feedback definition employed in quantum control theory and optics we call this type of feedback for ‘coherent feedback’ (see [9, 10] and references therein).

### Background compensation in a motif 2 based oscillatory homeostat

Fig 4a shows an example of a frequency-compensated oscillator, but now with the novel finding that it can also compensate for different but constant backgrounds.

**Fig 4.**
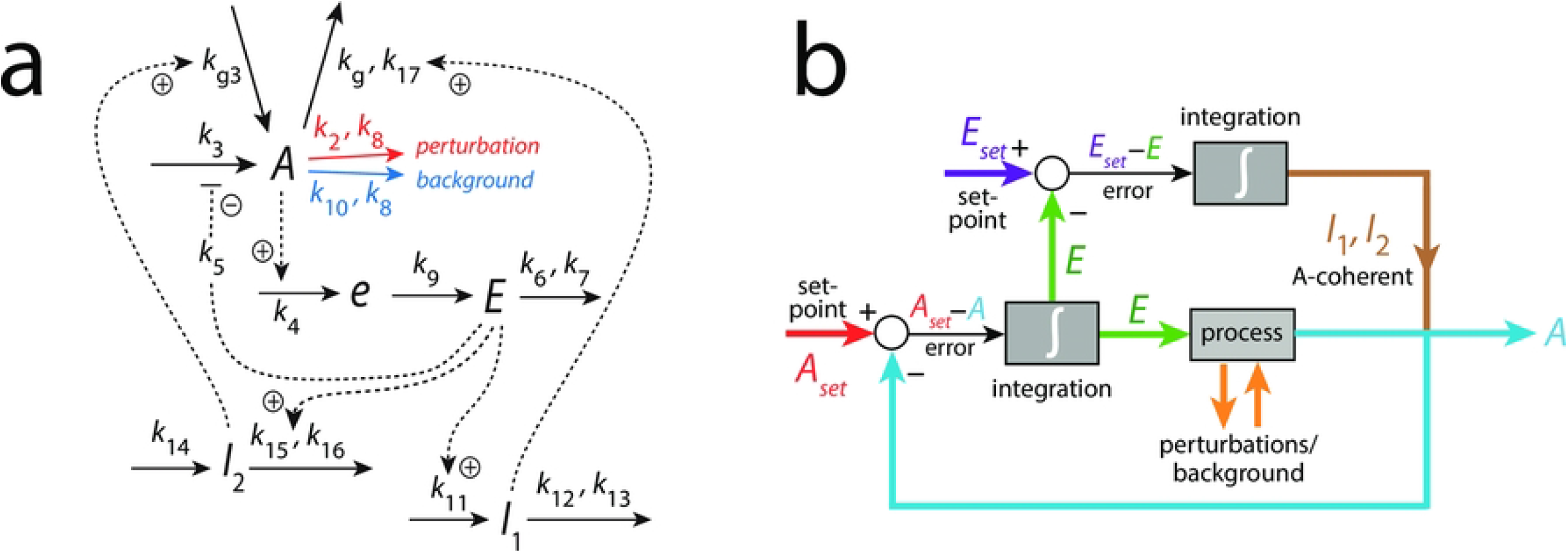
Frequency-compensated oscillator with background compensation by coherent feedback. Panel a: Reaction system based on derepression motif 2 (m2) [22] in the inner *A*-*e*-*E*-*A* negative feedback. Figs S12-S14 in the supporting information of Ref [11] describe some properties of this oscillator, but without having recognized at that time the ability to robustly compensate for backgrounds. Solid arrows indicate chemical reactions, while dashed lines show activations (plus signs) and one inhibition (minus sign). Panel b: Flow scheme indicating the additional control of *E* via *A* by controllers *I*_1_ and *I*_2_.

The center oscillator in Fig 4a is given by the *A*-*e*-*E*-*A* feedback loop based on derepression motif 2 [22], where *E* keeps *<A>* under homeostatic control (see rate equations and definitions of set-points below). Oscillations occur, because the removals of *A* and *E* are zero-order with respect to *A* and *E* and thereby construct a quasi-conservative oscillator. The intermediate *e* has been included to obtain limit-cycle oscillations [11]. *I*_1_ and *I*_2_ are controller species, which keep *<E>* under homeostatic control. It is the control of *<E>* by *I*_1_ and *I*_2_, which allows for the frequency homeostasis of the oscillator [11]. Their *A*-coherent feedback directly to *A* allows for robust background compensation. Since the central *A*-*e*-*E*-*A* negative feedback is an inflow controller it principally can only compensate for outflow perturbations [22]. The outflow perturbation considered here splits into two components: a constant (zero-order) background with rate constant *k*_10_ (Fig 4a outlined in blue) and a (zero-order) perturbation part where rate constant *k*_2_ undergoes a step-wise change (in Fig 4a outlined in red). Zero-order kinetics with respect to *A* are achieved by small *k*_8_ and *k*_17_ values, i.e. *A/*(*k*_8_+*A*) *≈* 1 and *A/*(*k*_17_+*A*) *≈* 1. Fig 4b shows the flow scheme and the control of *E* by *I*_1_ and *I*_2_ via the *A*-coherent part of the controller.

The rate equations are:

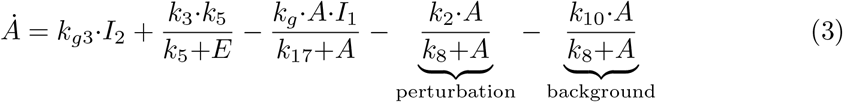

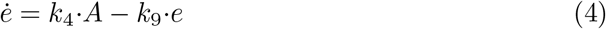

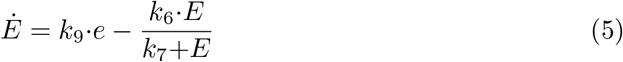

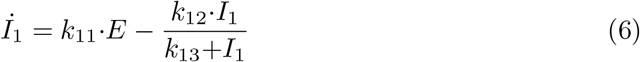

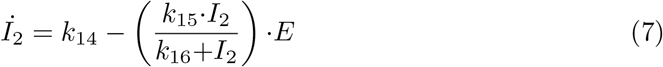

The set-point of *<A>* (*A*_*set*_) by controller *E* can be calculated from the steady state condition of the time averages:

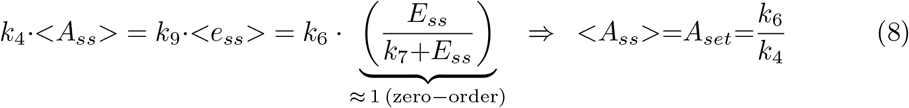

The zero-order condition with respect to *E* in Eq 8 ensures a robust perfect adaptation of *<A*_*ss*_*>* to *A*_*set*_ when the system oscillates, or of *A* to *A*_*set*_ in case the feedback loop is non-oscillatory [11]. Since the control of *A* by *E* is an inflow controller based on derepression of the flux *k*_3_*·k*_5_*/*(*k*_5_+*E*) the controller is active whenever *<A>* is below *A*_*set*_.

*E* is controlled by *I*_1_ and *I*_2_. They act as respectively outflow or inflow controllers [22] with respect to *<E>* (if oscillatory) or *E* (if non-oscillatory). Also here zero-order removals of both *I*_1_ and *I*_2_ ensure robust set-points. For controller *I*_1_ the steady state condition gives:

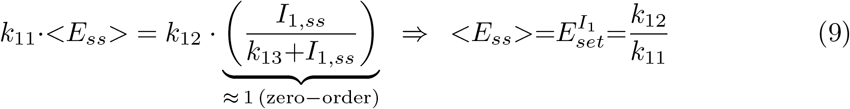

The *I*_1_ outflow controller becomes active whenever *<E>* is higher than 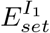

The set-point for the *I*_2_ inflow controller is determined by the steady state condition:

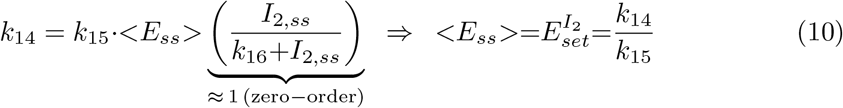

The *I*_2_ controller becomes active whenever *<E>* is lower than 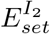. It should be noted that the values of the inflow/outflow set-points 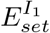 and 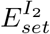 need to follow certain rules to guarantee that inflow and outflow controllers cooperate. In this case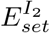 should be lower than 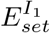, otherwise *I*_1_ and *I*_2_ will work against each other and windup will occur. For a discussion about windup in combined controllers, see Ref [22].

In the following we show how the above oscillator behaves in presence of a step-wise perturbation at different but constant backgrounds.

Fig 5 shows the oscillator’s behavior for a step-wise perturbation in *k*_2_ from 1.0 (phase 1) to 10.0 (phase 2) at a background *k*_10_=0.0. The time of change in *k*_2_ is indicated in each panel by a vertical arrow. Panel a shows the oscillations in *A* together with its average *<A>* (Eq 1), while panel b shows *E* and *<E>* (Eq 2). Panel c shows the changes in *I*_1_ and *I*_2_, and panel d shows the frequency (inverse of the period length). The resetting of the frequency to its pre-perturbation value is clearly seen. If *I*_1_ and *I*_2_ would not be present, *<A>* would be kept at *A*_*set*_=2.0 by a reduced (derepressed) *E* as seen in panel b at 100 time units. However, since *<E>* is also controlled by *I*_1_ and *I*_2_, i.e. between 5.0 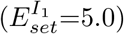 and 4.99 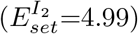*I*_1_ and *I*_2_ take over both for the control of *<A>* and of *<E>*.

**Fig 5.**
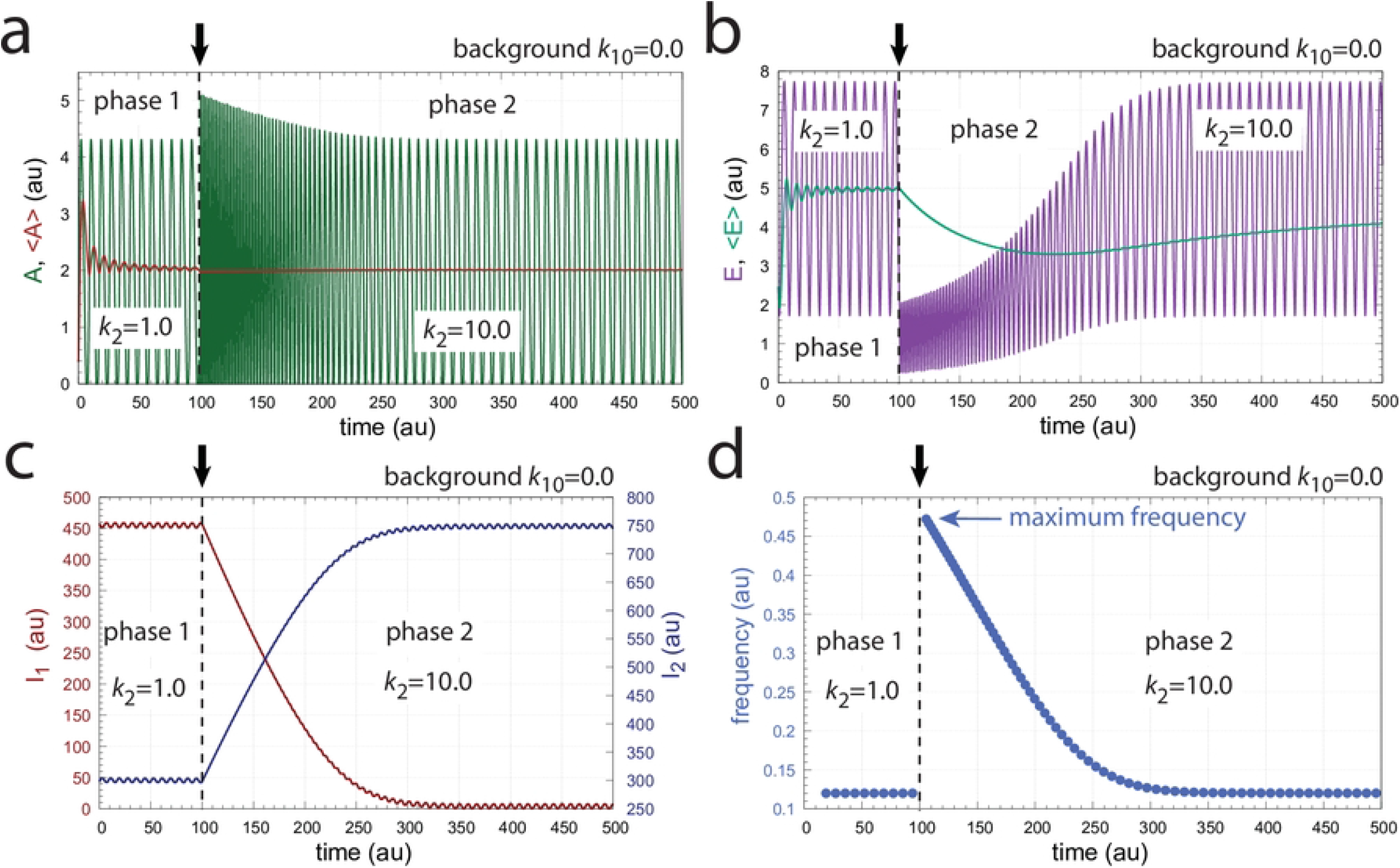
Frequency compensation of feedback scheme in Fig 4a for a step-wise change in *k*_2_ from 1.0 to 10.0 at a background of *k*_10_=0.0. Vertical arrows indicate the change in *k*_2_. Other rate constants: *k*_3_=100.0, *k*_4_=1.0, *k*_5_=0.1, *k*_6_=2.0, *k*_7_=*k*_8_=*k*_13_=*k*_16_=*k*_17_=1*×*10^*−*6^, *k*_9_=20.0, *k*_11_=1.0, *k*_12_=5.0, *k*_14_=4.99, *k*_15_=1.0, and *k*_*g*_=*k*_*g*3_=1*×*10^*−*2^. Initial concentrations: *A*_0_=0.3780, *E*_0_=2.4784, *e*_0_=1.5993*×*10^*−*2^, *I*_1,0_=4.5727*×*10^2^, *I*_2,0_=2.9817*×*10^2^ (see S1 Programs for python script).

Fig 6 shows the same perturbation in *k*_2_ as in Fig 5 but with a background of *k*_10_=2048.0. The increased removal of *A* by the background is compensated by an increase of *I*_2_ and a decrease of *I*_1_, which keep *<A>* and *<E>* at their respective set-points. The maximum frequency, which occurs directly after the *k*_2_ step is not affected. In other words, the sensitivity of the oscillator with respect to *k*_2_-step perturbations is compensated by *I*_1_ and *I*_2_ and is independent of the background *k*_10_.

**Fig 6.**
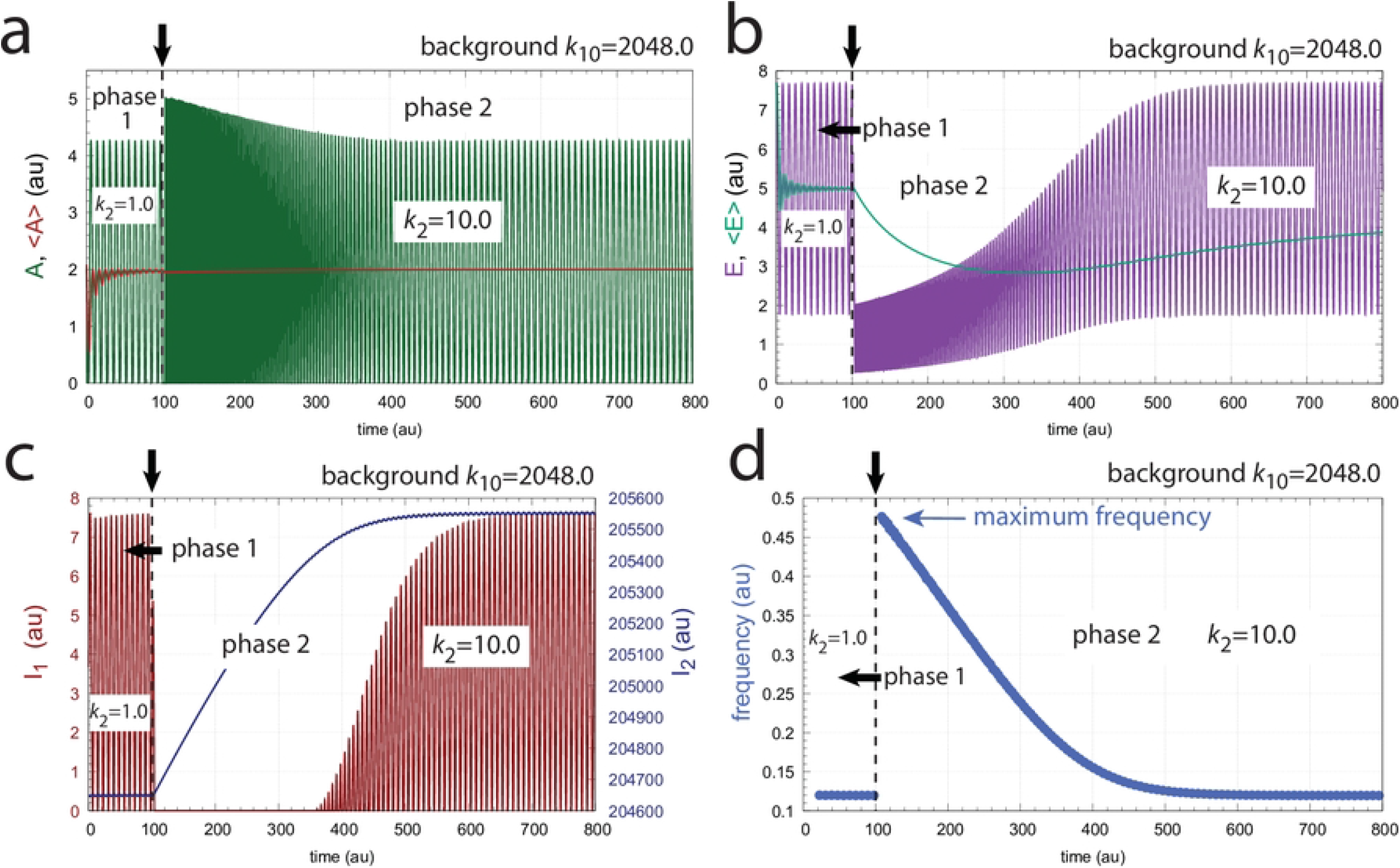
Frequency compensation of feedback scheme in Fig 4a for a step-wise change in *k*_2_ from 1.0 to 10.0 at a background of *k*_10_=2048.0. Vertical arrows indicate the change in *k*_2_. Other rate constants as in Fig 5. Initial concentrations: *A*_0_=2.1377, *E*_0_=7.6720, *e*_0_=1.0996*×*10^*−*1^, *I*_1,0_=3.4304, *I*_2,0_=2.0465*×*10^5^ (see S1 Programs for python script).

Fig 7 shows how the maximum frequency depends on *k*_2_ steps at different but constant backgrounds *k*_10_. The parallel lines indicate that the maximum frequency responses are independent of the background (“background compensation”).

**Fig 7.**
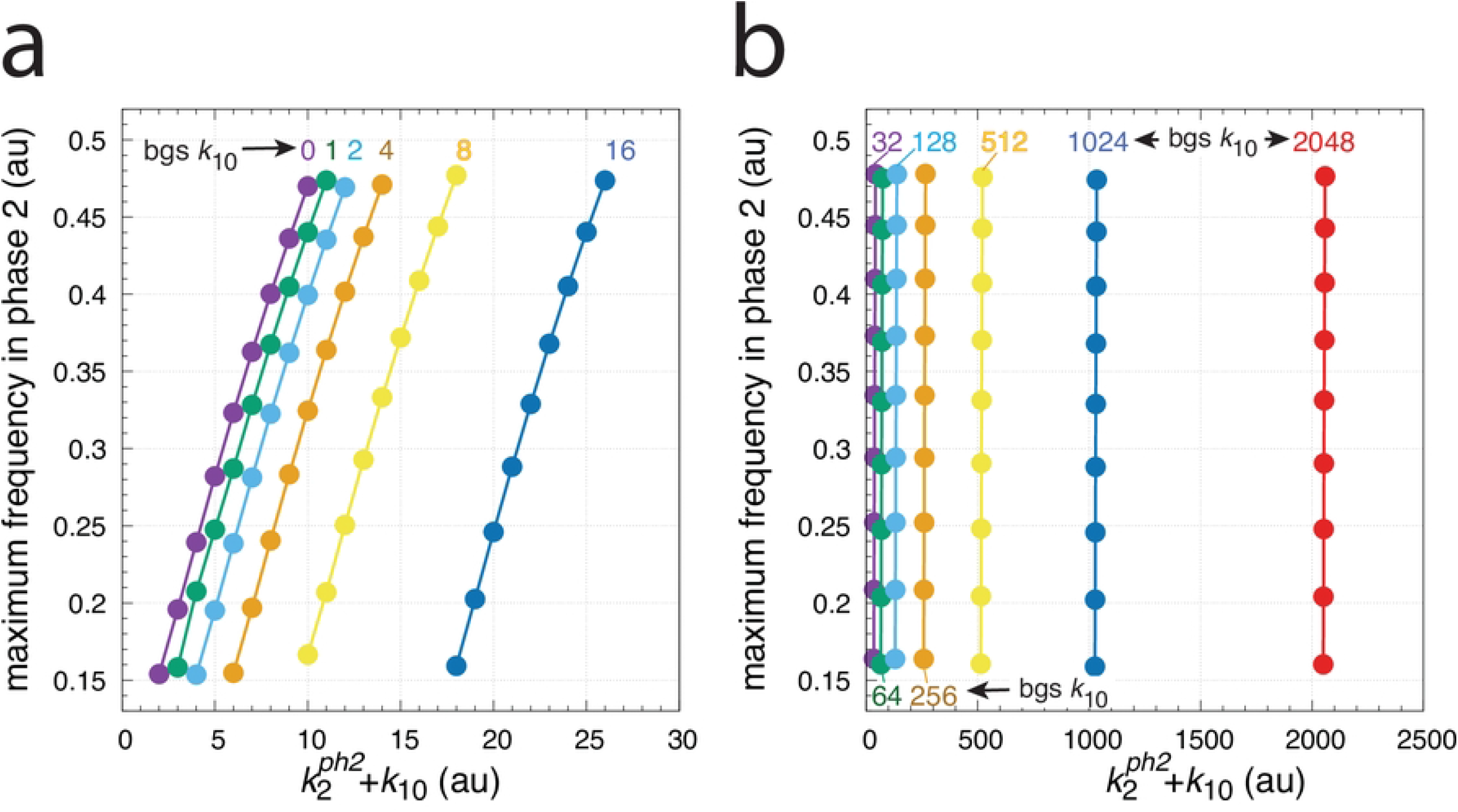
Background frequency compensation in the oscillator of Fig 4. The maximum frequency (see Fig 5d or Fig 6d) is plotted as a function of 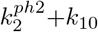 where 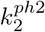is the *k*_2_ value during phase 2. The maximum frequency is determined for different step-wise *k*_2_ changes, i.e. for 1.0*→*2.0, 1.0*→*3.0,…, 1.0*→*9.0, up to 1.0*→*10.0, which occur at time t=100 at different but constant *k*_10_ backgrounds (bgs). Calculations have been performed analogous to Fig 5 and Fig 6. The *k*_10_ background values are 0, 1, 2, 4, up to 2048 (indicated in the figure). Other rate constants as in Fig 5. Initial concentrations: bgs 0-128, as in Fig 5; bg 256, *A*_0_=0.9866, *E*_0_=7.3508, *e*_0_=5.2447*×*10^*−*2^, *I*_1,0_=5.8243, *I*_2,0_=2.5447*×*10^4^; bg 512, *A*_0_=8.3872*×*10^*−*4^, *E*_0_=4.8793, *e*_0_=3.9572*×*10^*−*5^, *I*_1,0_=7.6544, *I*_2,0_=5.1046*×*10^4^; bg 1024, *A*_0_=1.7657, *E*_0_=7.6866, *e*_0_=9.1430*×*10^*−*2^, *I*_1,0_=4.2379, *I*_2,0_=1.0225*×*10^5^; bg 2048, as in Fig 6.

### Background compensation in a motif 8 (m8) based oscillatory homeostat

To provide an additional example of a frequency-compensated negative feedback oscillator with background compensation we use a m8 outflow control motif [22] for the center feedback. In this motif, *A* the controlled variable, inhibits the generation of the controller *E. E* on its side inhibits the removal of *A*. The outer controllers, *I*_1_ and *I*_2_, feed directly back to *A*. As for the m2 controller, oscillations in the central m8 oscillator are facilitated by removing *A* and *E* by zero-order processes. The scheme of this oscillator is shown in Fig 8. The rate equations are (‘pert’ stands for perturbation and ‘bg’ for background):

**Fig 8.**
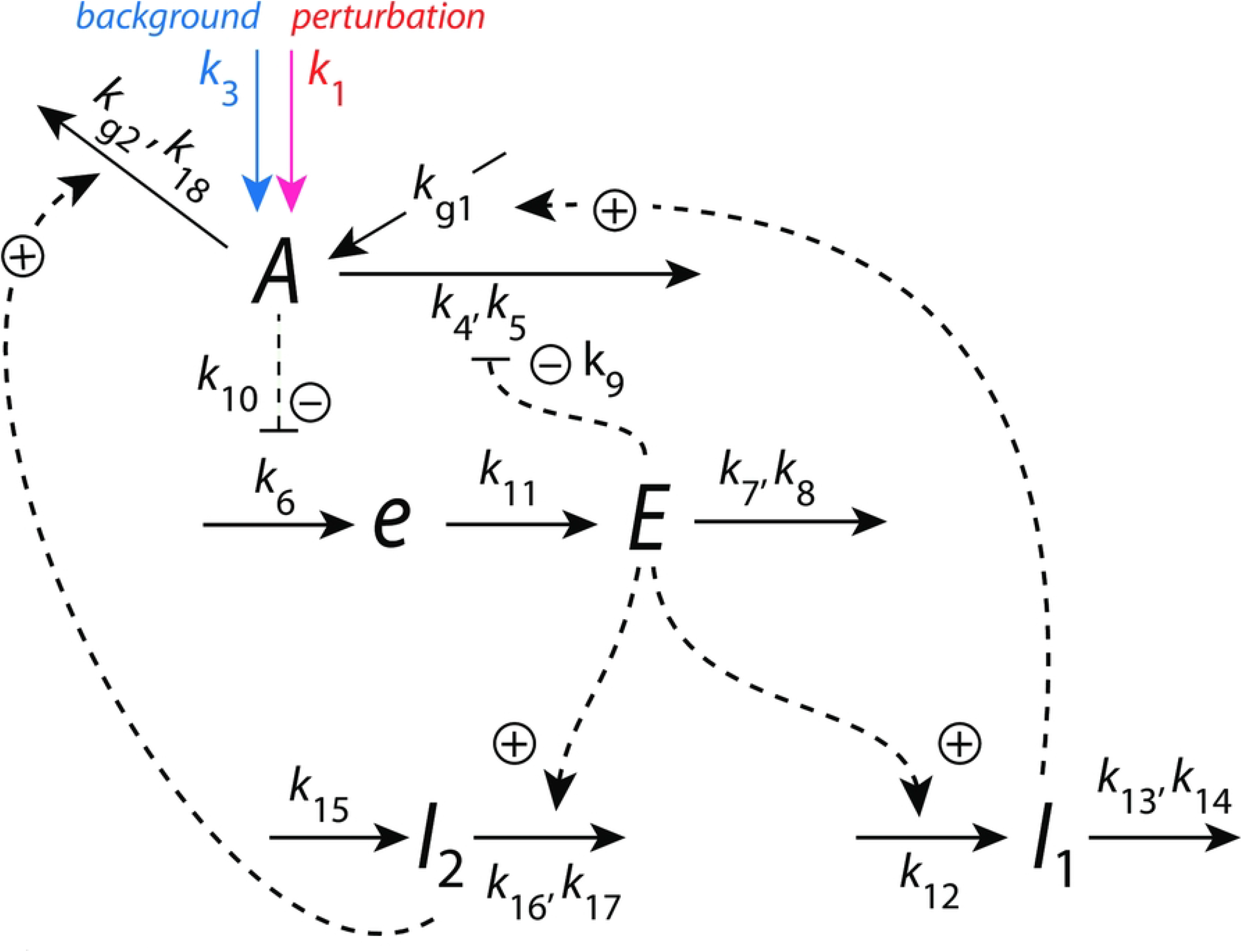
Frequency-compensated oscillator with background compensation by coherent feedback based on derepression motif m8 in the inner *A*-*e*-*E*-*A* negative feedback. Solid arrows indicate chemical reactions, while dashed lines show activations (plus signs) and inhibitions (minus sign).

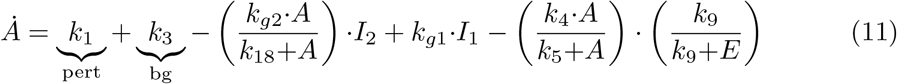

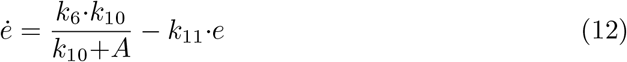

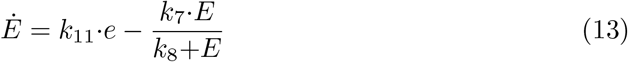

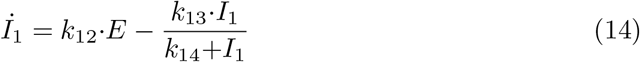

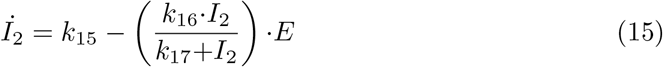

The inflow to *A* is divided into a step-wise perturbative component *k*_1_ (indicated in Eq 11 by ‘pert’ and outlined in red in Fig 8) and a background *k*_3_ (indicated in Eq 11 by ‘bg’ and outlined in blue in Fig 8). All other in- and outflows to and from *A* are compensatory fluxes.

As for the m2 oscillator above, we can calculate the set-points for *<E>* by *I*_1_ and *I*_2_ by setting Eqs 14 and 15 to zero and assume that *I*_1_ and *I*_2_ are removed by zero-order reactions, i.e.:

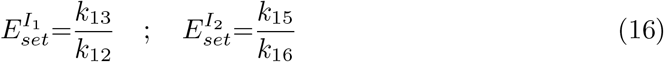

Unfortunately, for this scheme the oscillatory *A*_*set*_ cannot be calculated analytically. The closest analytical expression we can obtain is by setting Eqs 12 and 13 to zero, eliminating the *k*_11_*·e* term, and then calculating the time average of 1*/*(*k*_10_+*A*):

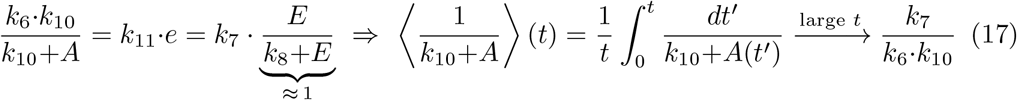

While calculations easily verify the right-hand side of Eq 17, *<A>* needs to be calculated numerically.

Fig 9 shows that the oscillator described in Fig 8 shows frequency homeostasis at different but constant *k*_3_ backgrounds. In panel a the background is *k*_3_=0.0, while in panel b we have *k*_3_=1024.0. In both cases the maximum frequency for a *k*_1_ step of 1.0*→*100.0 is the same, indicating that the maximum frequency is background compensated.

**Fig 9.**
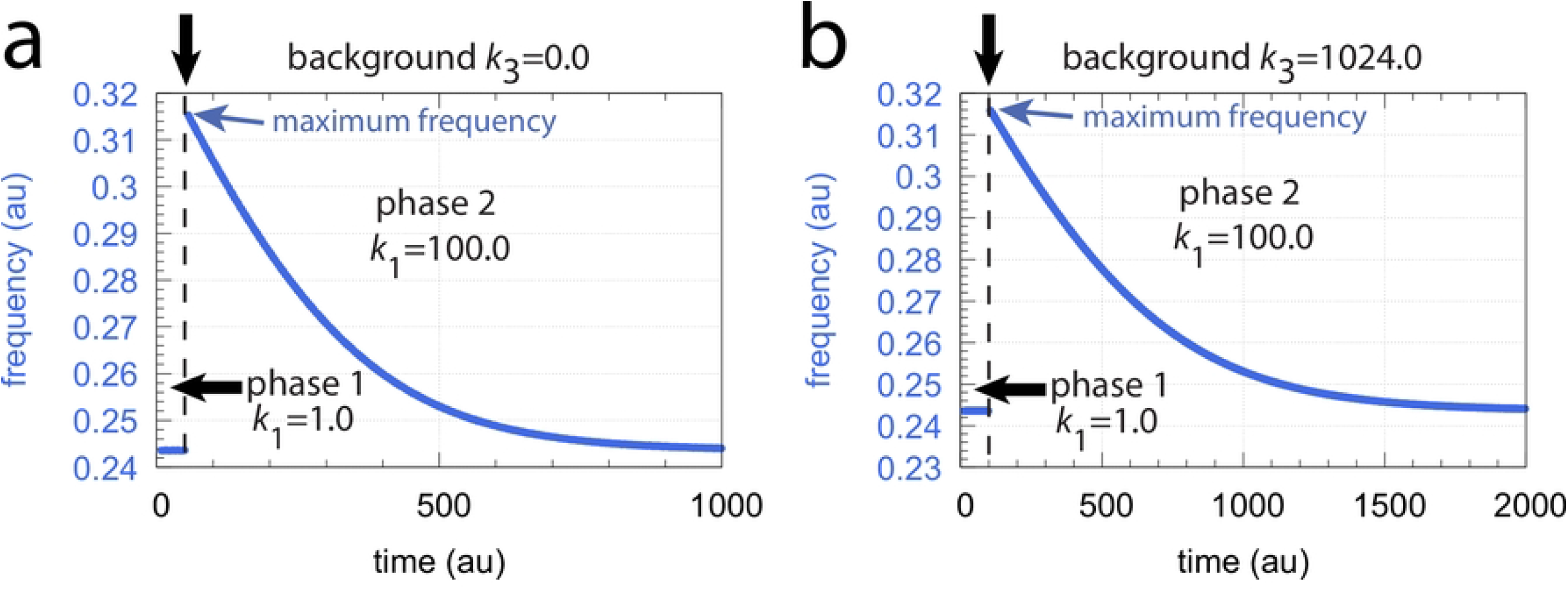
Frequency compensation/homeostasis in the oscillator described in Fig 8. In both panels a step-wise perturbation in *k*_1_ from 1.0 (phase 1) to 100.0 (phase 2) is applied (the step is indicated by the vertical arrows on top of the plots). In panel a, a constant background of *k*_3_=0.0 is applied (at both phases 1 and 2), while in panel b the background is 1024.0. Other rate constant values are: *k*_4_=1*×*10^4^, *k*_5_=*k*_8_=*k*_14_=*k*_17_=*k*_18_=1*×*10^*−*6^, *k*_6_=1*×*10^3^, *k*_7_=50.0, *k*_9_=0.1, *k*_11_=1.0, *k*_12_=5.0, *k*_13_=50.00, *k*_15_=50.0, *k*_16_=1.0 and *k*_*g*1_=*k*_*g*2_=1*×*10^*−*2^. Initial concentrations (*k*_3_=0.0): *A*_0_=3.3568*×*10^2^, *E*_0_=2.6209*×*10^1^, *e*_0_=7.3942, *I*_1,0_=2.4840*×*10^4^, *I*_2,0_=1.2768*×*10^4^. Initial concentrations (*k*_3_=1024.0): *A*_0_=3.6188, *E*_0_=1.8696*×*10^1^, *e*_0_=1.7115*×*10^2^, *I*_1,0_=4.6869, *I*_2,0_=9.0420*×*10^4^. Two python scripts, which in addition show the variations of *A, E, I*_1_, and *I*_2_, are included in S1 Programs.

Fig 10 shows maximum frequencies for different *k*_1_ steps and *k*_3_ backgrounds. There, different but constant *k*_3_ backgrounds are applied with values 0, 2, 4, 8, 16, 32, 64, 128, 256, 512, and 1024. Step perturbations in *k*_1_ are applied starting with a 1.0 (phase 1) to 10.0 (phase 2) step and ends with a 1.0 (phase 1) to 500.0 (phase 2) step by successively increasing the *k*_1_ values in phase 2 by 10.0. As the parallel lines in Fig 10 show, once the oscillator has reached steady state for a certain background, the maximum frequencies, although dependent on the *k*_1_ step become independent of the background *k*_3_.

**Fig 10.**
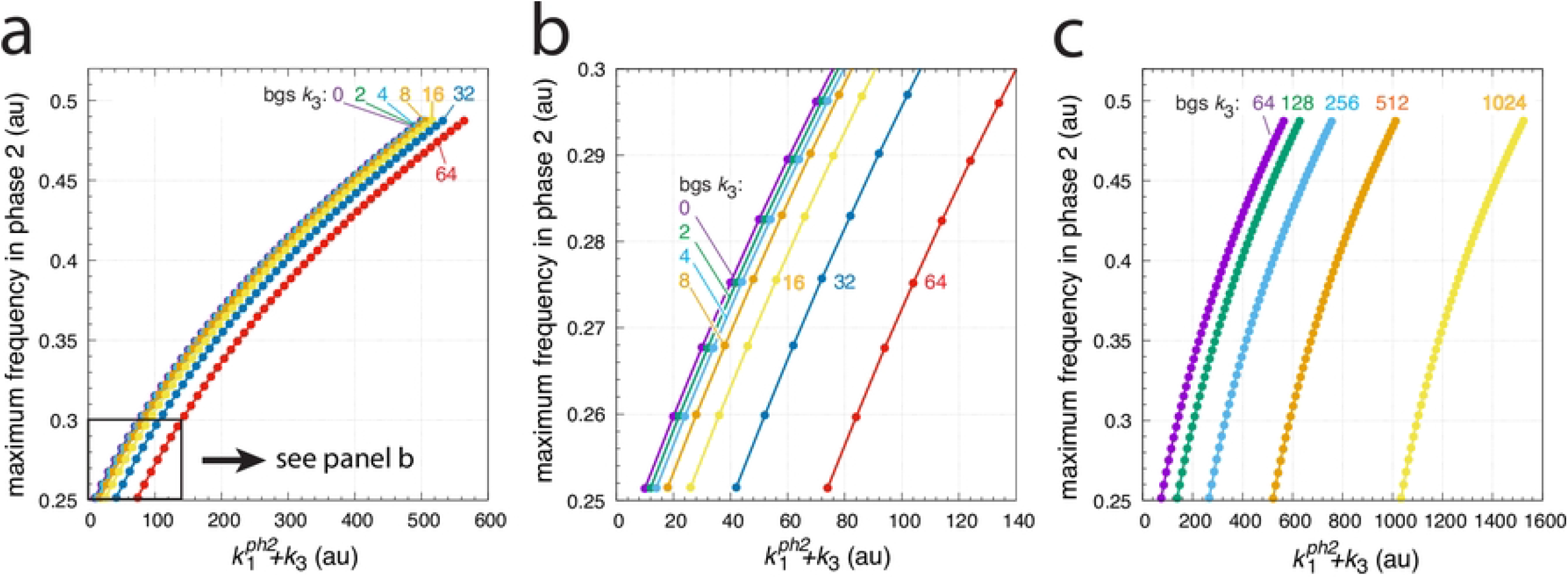
Background (bg) frequency compensation in the oscillator of Fig 8. The maximum frequency (see Fig 9) is plotted as a function of the sum of 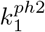 and background *k*_3_, where 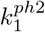 is the *k*_1_ value in phase 2. In analogy with the calculations in Fig 5 and Fig 6 the maximum frequency is determined for different step-wise *k*_1_ changes, i.e. for 1.0*→*10.0, 1.0*→*20.0,…, 1.0*→*30.0, up to 1.0*→*500.0, which occur at time t=100 at different but constant *k*_3_ backgrounds (bgs). The *k*_3_ background values are 0, 2, 4, up to 1024 (indicated in the figure). Other rate constants as in Fig 9. Initial concentrations: bg 0: as in Fig 9a; bg 2: *A*_0_=3.4008*×*10^2^, *E*_0_=2.3906*×*10^1^, *e*_0_=7.0224, *I*_1,0_=2.4739*×*10^4^, *I*_2,0_=1.2869*×*10^4^; bg 4: *A*_0_=3.4461*×*10^2^, *E*_0_=2.1393*×*10^1^, *e*_0_=6.6417, *I*_1,0_=2.4637*×*10^4^, *I*_2,0_=1.2971*×*10^4^; bg 8: *A*_0_=4.8073*×*10^2^, *E*_0_=6.0165*×*10^1^, *e*_0_=1.0914*×*10^2^, *I*_1,0_=2.4401*×*10^4^, *I*_2,0_=1.3207*×*10^4^; bg 16: *A*_0_=4.3570, *E*_0_=1.9953*×*10^1^, *e*_0_=1.6964*×*10^2^, *I*_1,0_=2.4005*×*10^4^, *I*_2,0_=1.3603*×*10^4^; bg 32: *A*_0_=3.9151*×*10^1^, *E*_0_=5.4663*×*10^1^, *e*_0_=1.1875*×*10^2^, *I*_1,0_=2.3201*×*10^4^, *I*_2,0_=1.4407*×*10^4^; bg 64: *A*_0_=3.0270*×*10^2^, *E*_0_=4.1000*×*10^1^, *e*_0_=1.0456*×*10^1^, *I*_1,0_=2.1646*×*10^4^, *I*_2,0_=1.5962*×*10^4^; bg 128: *A*_0_=3.2021*×*10^2^, *E*_0_=3.3534*×*10^1^, *e*_0_=8.7470, *I*_1,0_=1.8443*×*10^4^, *I*_2,0_=1.9165*×*10^4^; bg 256: *A*_0_=6.4511*×*10^1^, *E*_0_=6.8158*×*10^1^, *e*_0_=9.3623*×*10^1^, *I*_1,0_=1.2002*×*10^4^, *I*_2,0_=2.5606*×*10^4^; bg 512: *A*_0_=2.6297*×*10^2^, *E*_0_=5.5584*×*10^1^, *e*_0_=1.5294*×*10^1^, *I*_1,0_=3.2525*×*10^3^, *I*_2,0_=4.2375*×*10^4^; bg 1024: as in Fig 9b. Panel a shows an overview of the maximum frequencies up to background 64, while panel b shows a blown-up part indicated in panel a. Panel c shows the maximum frequencies for backgrounds from 64 up to 1024.

**Fig 11.**
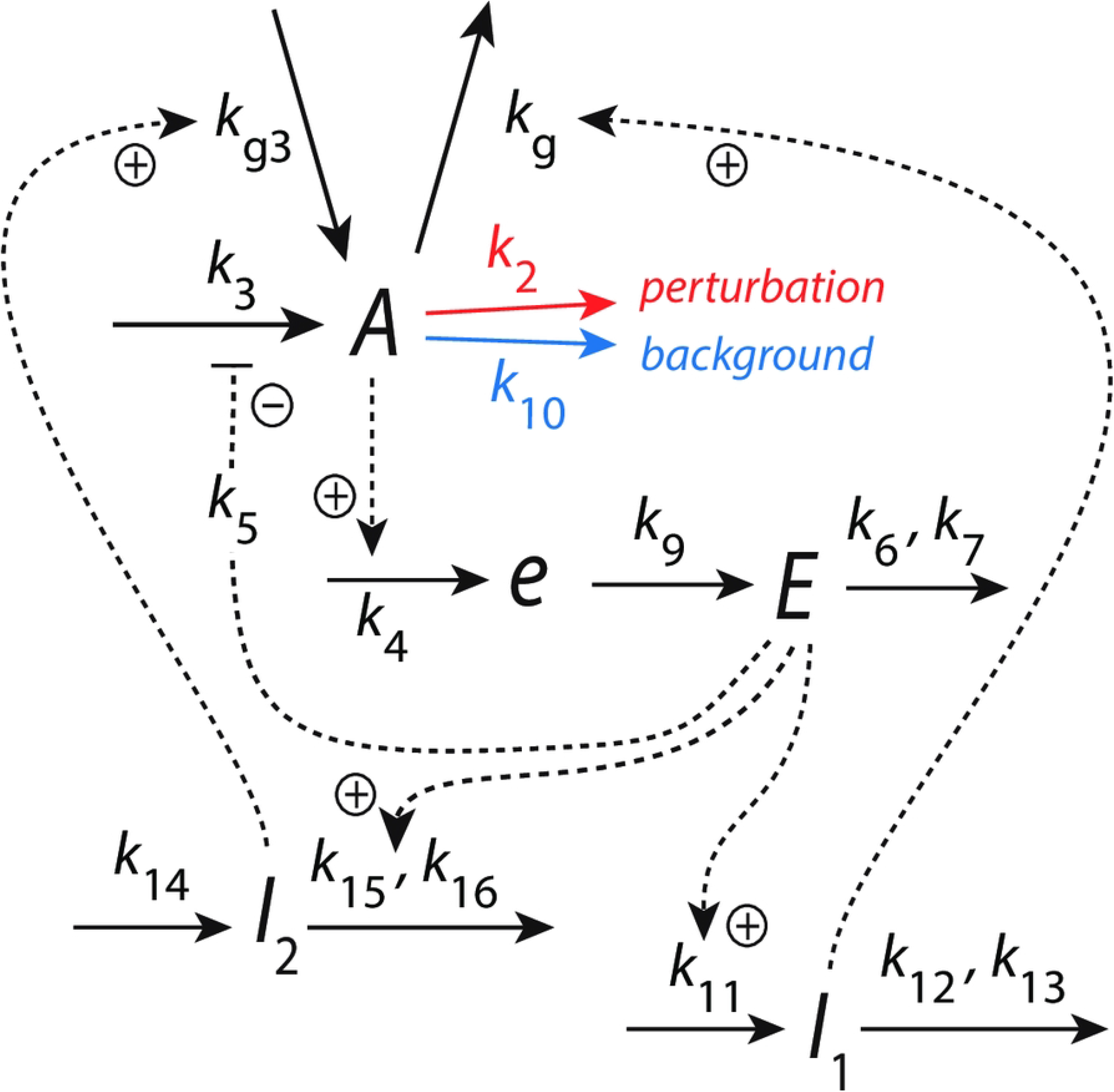
Same scheme as Fig 4a, but to facilitate a non-oscillatory homeostat all *A*-removing reactions are changed to first-order kinetics with respect to *A*.

### Background compensation in non-oscillatory homeostats

In this section we look at background compensation in non-oscillatory homeostats where *E* is controlled by *I*_1_ and *I*_2_ via coherent feedback. We show two examples: in the first one the controller’s response after a step perturbation is significantly damped, while in the other example the response shows a larger train of (damped) oscillations. In both cases the response profiles of the controlled variables *A* and *E* are preserved, independent of the background.

For the first example we use the oscillator scheme from Fig 4a. To go over to a non-oscillatory mode, we change the kinetics for all *A*-removing reactions from zero-order to first-order kinetics with respect to *A*. The rate equation of *A* becomes (compare with Eq 3):

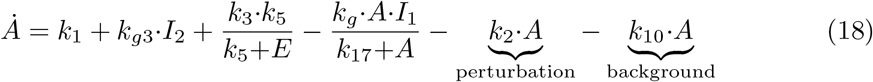

while the rate equations for the other components (Eqs 4-7) remain the same.

Fig 12 gives an overview of the results. In panel a the maximum excursions Δ*A* after the step (see inset) are plotted for different backgrounds *k*_10_ as a function of the sum of the phase 2 *k*_2_ value and *k*_10_. For each background, the nine *k*_2_ steps 1*→*2, 1*→*3, ….,1*→*9, and 1*→*10 are applied and Δ*A* is determined. The parallel lines indicate robust background compensation, i.e. Δ*A* is the same for a certain defined step, independent of the background. Panel b shows the situation for a *k*_2_ 1*→* 10 step when background *k*_10_=0. In panel c the same step is applied, but the background has been increased to *k*_10_=10. Comparing Figs 12b and 12c shows that profiles in both *A* and *E* are the same with *A*_*set*_=2.0 and *E*_*set*_=100.

**Fig 12.**
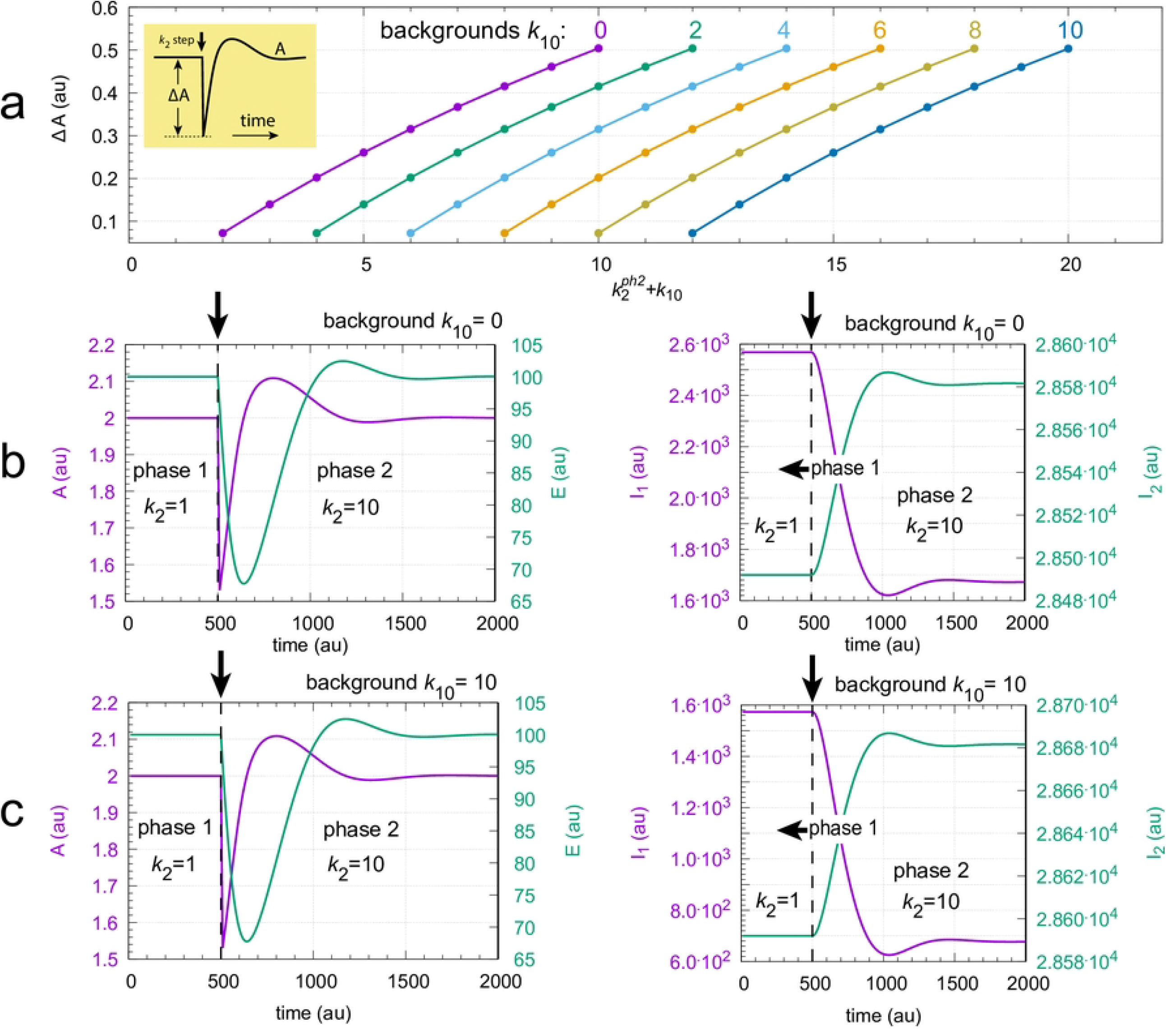
Background compensation in the non-oscillatory feedback scheme of Fig 11. Panel a: Each colored curve shows the values of Δ*A* for the nine *k*_2_ steps: 1*→*2, 1*→*3,…,1*→*10 at *k*_10_ background levels: 0, 2, 4,…,8, 10. Inset shows how Δ*A* is defined. 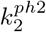 is the value of *k*_2_ during phase 2. Panels in b: Time profiles of *A, E* (left panel) and *I*_1_, *I*_2_ (right panel) for a 1*→*10 *k*_2_ step at background *k*_10_=0.0. The change in *k*_2_ is applied at time t=500 indicated by the vertical arrow. Panels in c are similar to the panels in b with the difference that background *k*_10_ is 10.0. Other rate constants: *k*_3_=5*×*10^3^, *k*_4_=1.0, *k*_5_=0.5, *k*_6_=2.0, *k*_7_=1*×*10^*−*5^, *k*_9_=2.0, *k*_11_=0.1, *k*_12_=10.0, *k*_13_=*k*_16_=1*×*10^*−*4^, *k*_14_=1.0, *k*_15_=0.01, *k*_*g*_=0.01, *k*_*g*3_=1*×*10^*−*3^. Initial concentrations for panel b: *A*_0_=2.0, *E*_0_=100.0, *e*_0_=1.0, *I*_1,0_=2.5684*×*10^3^, *I*_2,0_=2.8492*×*10^4^. Initial concentrations for panel c: *A*_0_=2.0, *E*_0_=100.0, *e*_0_=1.0, *I*_1,0_=1.5734*×*10^3^, *I*_2,0_=2.8592*×*10^4^. For python scripts showing the results of panels b and c, please see Supporting information S1 Programs.

Fig 13 shows another example of coherent feedback. Here we have two inflow controllers *E*_1_ and *E*_2_, but only *E*_2_ is connected to *A* via *I*_1_ and *I*_2_ through a coherent feedback. The reason why we looked at two *E*-controllers was to see whether *E*_2_ alone, i.e. without the help of *I*_1_ and *I*_2_, was able to compensate backgrounds. This, however, turned out not to be the case and *I*_1_ and *I*_2_ were included to control *E*_2_.

**Fig 13.**
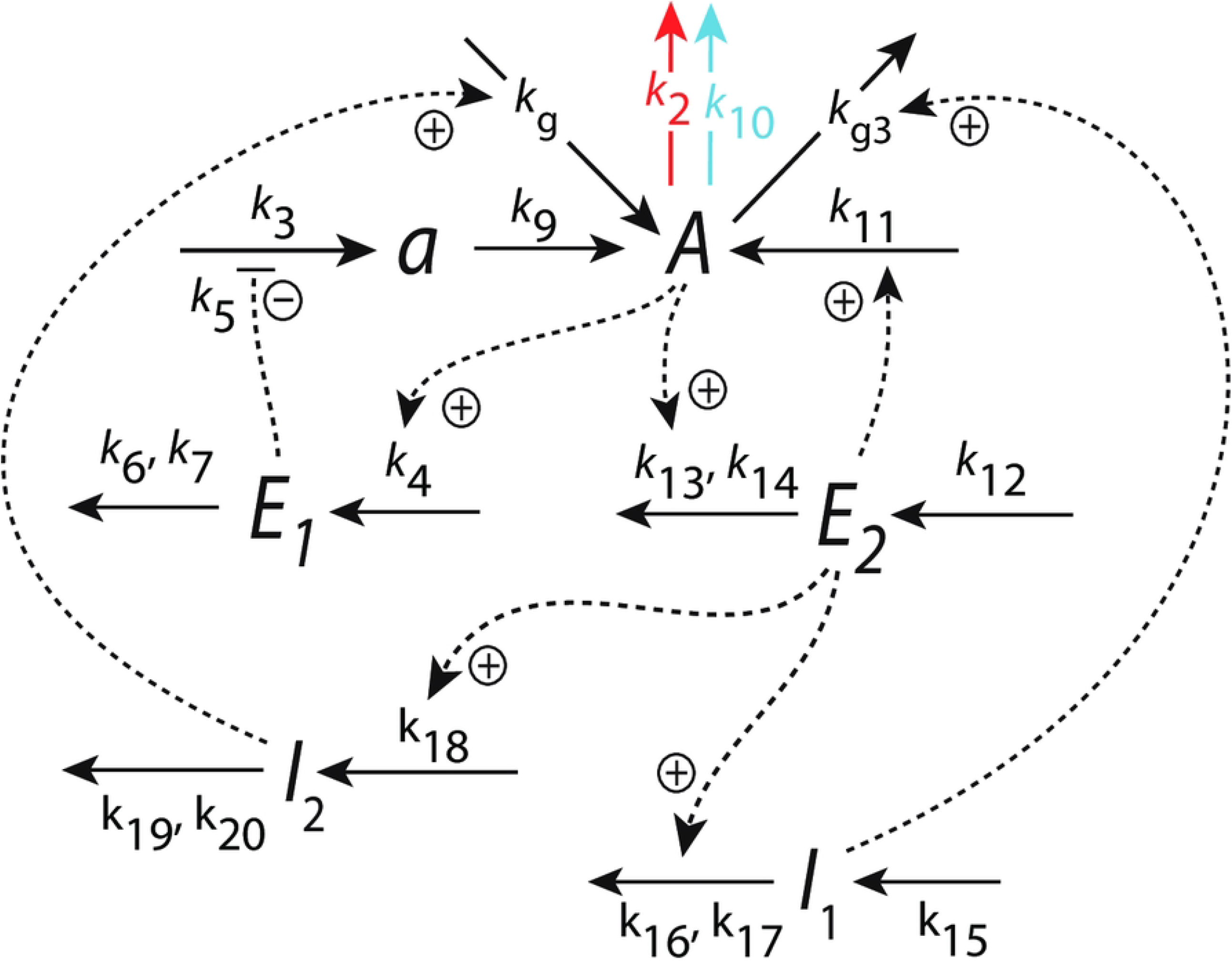
Coherent feedback loop *A*-*E*_2_-(*I*_1_,*I*_2_)-A with an additional inflow control of *A* by *E*_1_.

The rate equations are:

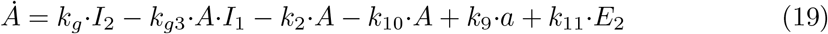

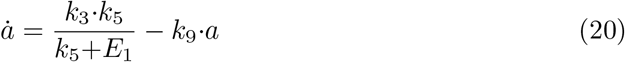

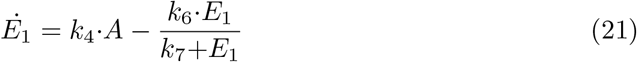

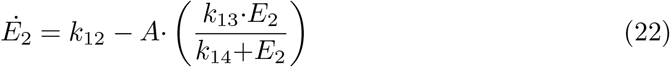

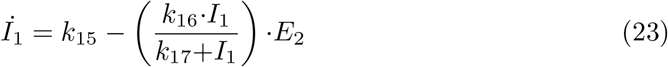

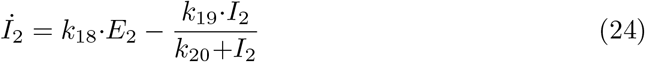

*E*_1_ and *E*_2_ provide two set-points for *A*: one, 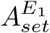, by setting Eq 21 to zero and solving for the steady state level of *A* under zero-order conditions, and the other, 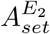 by doing the same for Eq 22. This gives:

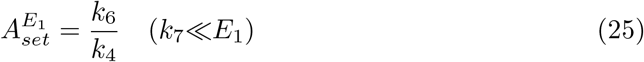

and

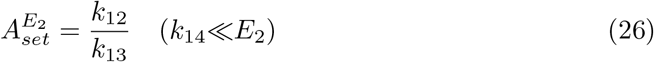

In the calculations we have set 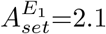 and 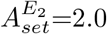 Since the *E*_2_ inflow controller has a lower set-point in comparison with *E*_1_, *E*_2_ will take over the control of *A* [22], while *E*_1_ will be inactive and allow a constant inflow to *A* via *a*.

Fig 14 shows that in this system a 1*→*10 perturbation in *k*_2_ induces a train of damped oscillations with background (*k*_10_) independent concentration profiles. In panels a and b Δ*A* (for definition see inset in Fig 12) is shown as a function of increasing *k*_2_ steps at different but constant *k*_10_ backgrounds. Panel c shows the time profile in *A* for a 1*→*10 *k*_2_ step with a *k*_10_ background of 0. In panel d the same step is applied but now with a background of *k*_10_=1024. One clearly sees the conserved background-independent transition profiles in *A*.

**Fig 14.**
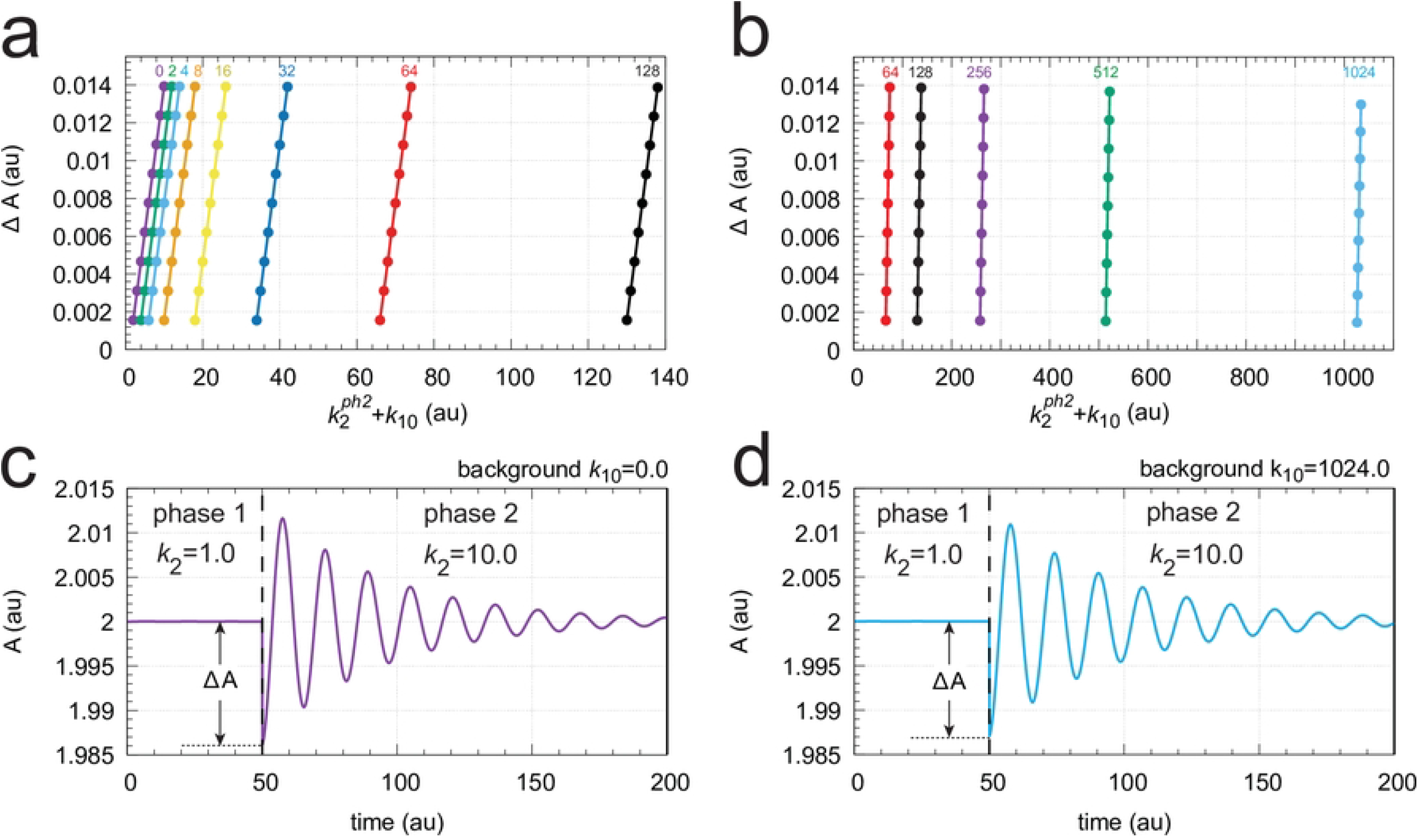
Background compensation by coherent feedback in the scheme of Fig 13. Panels a and b: Δ*A* as a function of 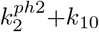 for different but constant backgrounds. 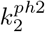is the *k*_2_ value in phase 2. The background values (*k*_10_) are indicated above the colored curves. Panels c and d: Concentration profiles in *A* when a *k*_2_ 1*→*10 step is applied at respective backgrounds of *k*_10_=0.0 and 1024.0. Other rate constants: *k*_3_=2.5*×*10^3^, *k*_4_=1.0, *k*_5_=0.1, *k*_6_=2.1, *k*_7_=1*×*10^*−*5^, *k*_9_=0.5, *k*_11_=0.5, *k*_12_=200.0, *k*_13_=100, *k*_14_=*k*_17_=*k*_20_=1*×*10^*−*5^, *k*_15_=1*×*10^3^, *k*_16_=10.0, *k*_18_=1.0, *k*_19_=99.99, *k*_*g*_=*k*_*g*3_=0.1. Initial concentrations for c: *A*_0_=2.0, *E*_1,0_=2.0*×*10^*−*4^, *E*_2,0_=100.0, *a*_0_=4.99*×*10^3^, *I*_1,0_=2.5684*×*10^3^, *I*_2,0_=2.8492*×*10^4^. Initial concentrations for d: *A*_0_=2.0, *E*_1,0_=2.0*×*10^*−*4^, *E*_2,0_=100.0, *a*_0_=4.99*×*10^3^, *I*_1,0_=3.5195*×*10^3^, *I*_2,0_=2.0888*×*10^3^. Supporting information S1 Programs includes the python scripts showing the results of panels c and d.

### Frequency homeostasis without background compensation

Fig 15a shows an oscillator scheme which we described previously in relation to robust frequency homeostasis [11]. We wondered whether frequency homeostasis would imply background compensation, but found out that this is not the case. In this case *I*_1_, *I*_2_, and *E* do not feed back coherently to *A*, but (incoherently) to *a*, which is a precursor of *A*.

**Fig 15.**
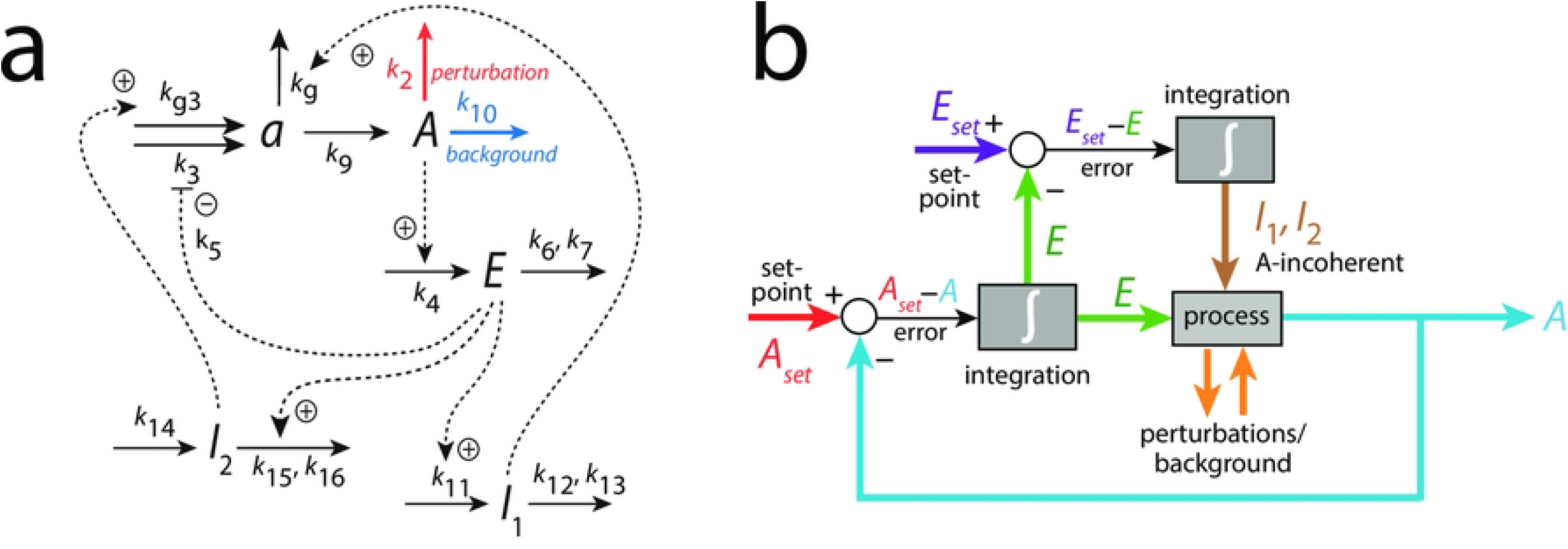
Oscillator based on motif 2 [11, 22] with *A*-incoherent feedback, where *E, I*_1_, and *I*_2_ feed back to *a*, a precursor of *A*. Panel a: reaction scheme. Panel b: Flow scheme. For rate equations, see main text.

The rate equations are (‘pert’ stands for perturbation and ‘bg’ for background):

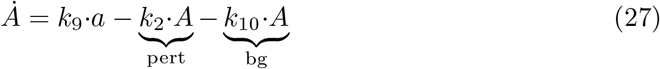

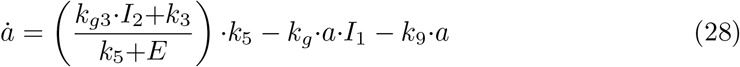

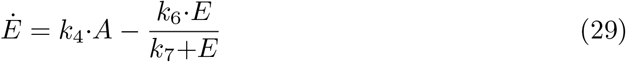

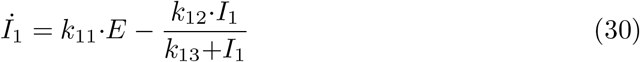

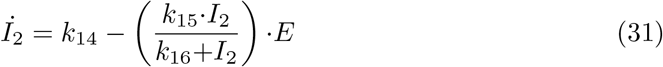

Fig 16 shows an example of oscillations for the scheme in Fig 15 with background *k*_10_=0 and a step perturbation in *k*_2_ from 1 (phase 1) to 10 (phase 2). It may be noted that in this case oscillations occur although the removal reactions of *a* and *A* are first-order with respect to *a* and *A*, indicating that first-order processes are only a ‘weak’ condition to abolish oscillatory behavior, as has been indicated in the above section ‘Background compensation in non-oscillatory homeostats’.

**Fig 16.**
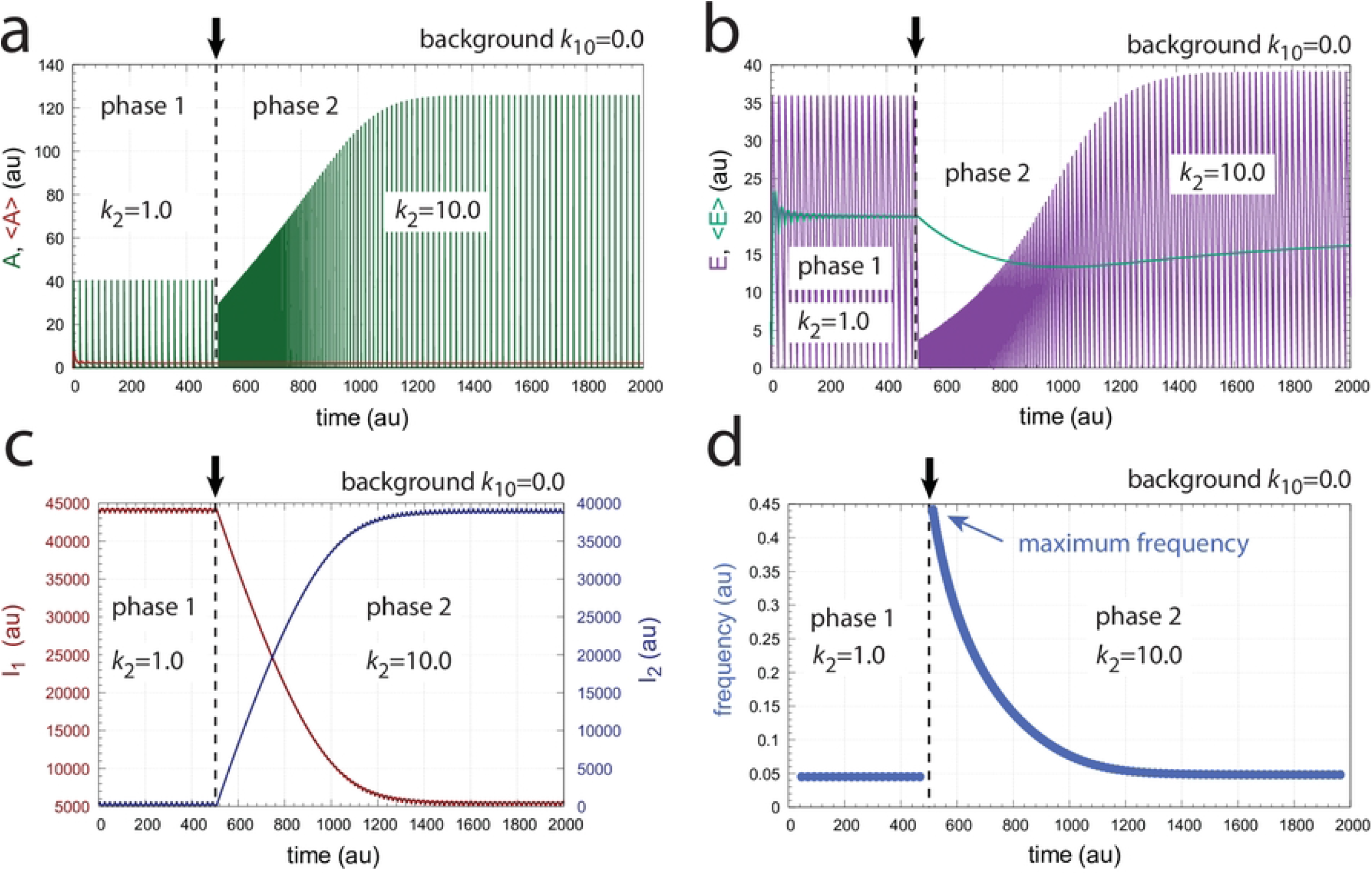
Frequency homeostasis in the oscillator of Fig 15. Background *k*_10_=0.0. A *k*_2_ step 1*→*10 occurs at time t=500 indicated by the vertical arrows. Panel a: Concentration of *A* and average *<A>* as a function of time. Panel b: Concentration of *E* and average *<E>* as a function of time. Panel c: Concentrations of *I*_1_ and *I*_2_ as a function of time. Panel d: Frequency as a function of time. Other rate constants: *k*_3_=1*×*10^6^, *k*_4_=1.0, *k*_5_=1*×*10^*−*6^, *k*_6_=2.0, *k*_7_=*k*_13_=*k*_16_=1*×*10^*−*6^, *k*_9_=2.0, *k*_11_=5.0, *k*_12_=100.0, *k*_14_=99.99, *k*_15_=5.0, *k*_*g*_=1*×*10^*−*3^, and *k*_*g*3_=100.0. Initial concentrations: *A*_0_=5.6920*×*10^*−*3^, *E*_0_=6.1163, *a*_0_=3.6221*×*10^*−*3^, *I*_1,0_=4.4051*×*10^4^, *I*_2,0_=2.7566*×*10^2^. See S1 Programs for python scripts.

Fig 16 clearly shows the occurrence of frequency homeostasis. However, when the oscillator is tested for different but constant *k*_10_ backgrounds with changed *k*_2_ steps the maximum frequency decreases with increasing backgrounds. Fig 17a shows the decrease of the maximum frequency and loss of robust background compensation at four different *k*_10_ backgrounds when *k*_2_ steps are applied from 1*→*2 up to 1*→*10 in analogy to the calculation shown in Fig 16. When using a logarithmic ordinate (Fig 17b) lines appear more or less parallel, which may give the illusion that the system responds in a background compensated way.

**Fig 17.**
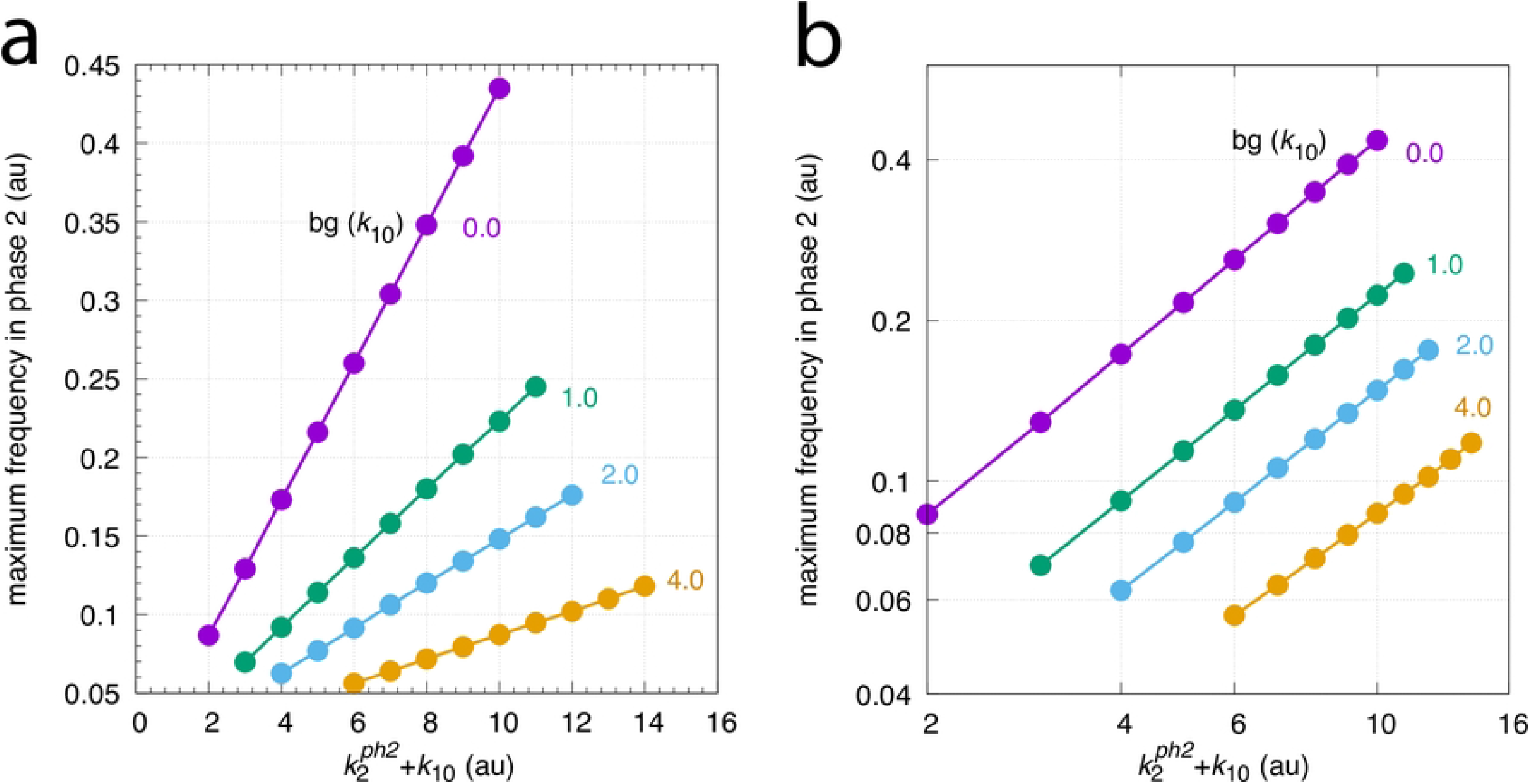
Maximum frequencies as a function of 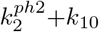 where 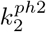 is the *k*_2_ value in phase 2. Calculations were performed with rate constants as described in Fig 16. Panel a show results with linear scaling of axes, while panel b shows the same data set as double-logarithmic plots. Initial concentrations: bg (*k*_10_)=0.0, see legend of Fig 16; bg (*k*_10_)=1.0, *A*_0_=2.5946*×*10^*−*3^, *E*_0_=25.4830, *a*_0_=2.6844*×*10^*−*3^, *I*_1,0_=3.0980*×*10^4^, *I*_2,0_=1.3296*×*10^4^; bg (*k*_10_)=2.0, *A*_0_=5.0041*×*10^*−*3^, *E*_0_=15.8930, *a*_0_=7.8102*×*10^*−*3^, *I*_1,0_=2.2995*×*10^4^, *I*_2,0_=2.1181*×*10^4^; bg (*k*_10_)=4.0, *A*_0_=4.7328*×*10^*−*3^, *E*_0_=21.6050, *a*_0_=1.2043*×*10^*−*2^, *I*_1,0_=1.3516*×*10^4^, *I*_2,0_=3.0610*×*10^4^.

## Is retinal light adaptation background compensated?

Based on the comment in Ref [2] that the parallel lines in Fig 2 indicate the same response at different backgrounds and involve a form of compensation mechanism, we became interested to look into the conditions how background compensation could occur. This requires of course how the term ‘background compensation’ is defined. We have applied the following, we think rather intuitive definition, where background compensation for a negative feedback system means the presence of a compensatory mechanism, which, when a perturbation is applied, the same response in a controlled variable occurs *independent* of an applied constant background in relationship to the perturbation. This definition is, however, not in agreement with the results shown in Fig 2, for example for the backgrounds indicated by the red, blue and green curves. Increasing the background from the red, blue to the green curve leads to a reduction in the average maximum frequency when a test spot luminance of 1*×*10^*−*2^ cd/m^2^ is applied, as indicated by the vertical dashed bar. In fact, the adaptation behavior shown in Fig 1 can show an analogous behavior as in Fig 2.

To see this we use the model where the response amplitude *V* of retinal cells with respect to a light perturbation *I* are described by a Hill-type Michaelis-Menten equation of the form

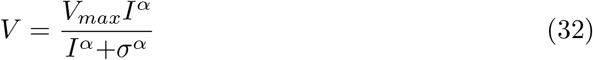

The cooperativity *α* is 1.0 for photoreceptor cells, but found to be 0.7 to 0.8 for horizontal cells, 1.2-1.4 for bipolar and sustained ganglion cells, and about 3.4 for transient ganglion cells (for an overview see [3]).

We consider here the response kinetics of rods and cones, i.e. *σ*=1 with

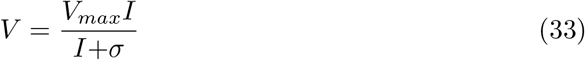

As pointed out by Naka and Rushton [32], in the presence of a background *I*_0_ the response *V*_1_ upon a perturbation *I*_1_ of a single pigment system will follow Eq 33, but with an increase of *s* to *s*_1_=*s*+*I*_0_ and a scaling of *V*_*max*_ by a factor of *s*/(*s*+*I*_0_). This can be shown as follows:

In the presence of a constant background *I*_0_ Eq 33 gives

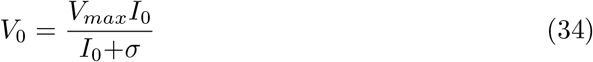

If a light perturbation *I*_1_ is applied in addition to background *I*_0_ the total response amplitude is

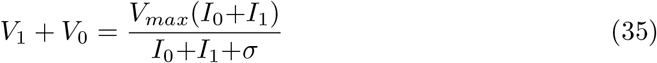

Subtracting Eq 34 from Eq 35 gives

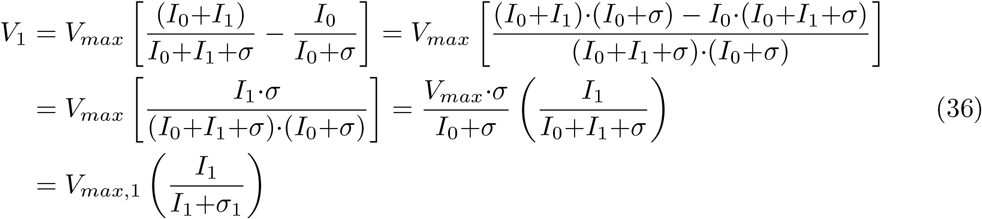

Fig 18 shows Eq 33 with six different *σ* values which mimick six different background levels. For the sake of simplicity we have set *V*_*max*_=1. In panel a both axes are linear, while in panel b the ordinate is logarithmic and the abscissa is linear. In panel c the ordinate is linear and the abscissa is logarithmic. Finally, in panel d both axes are logarithmic.

**Fig 18.**
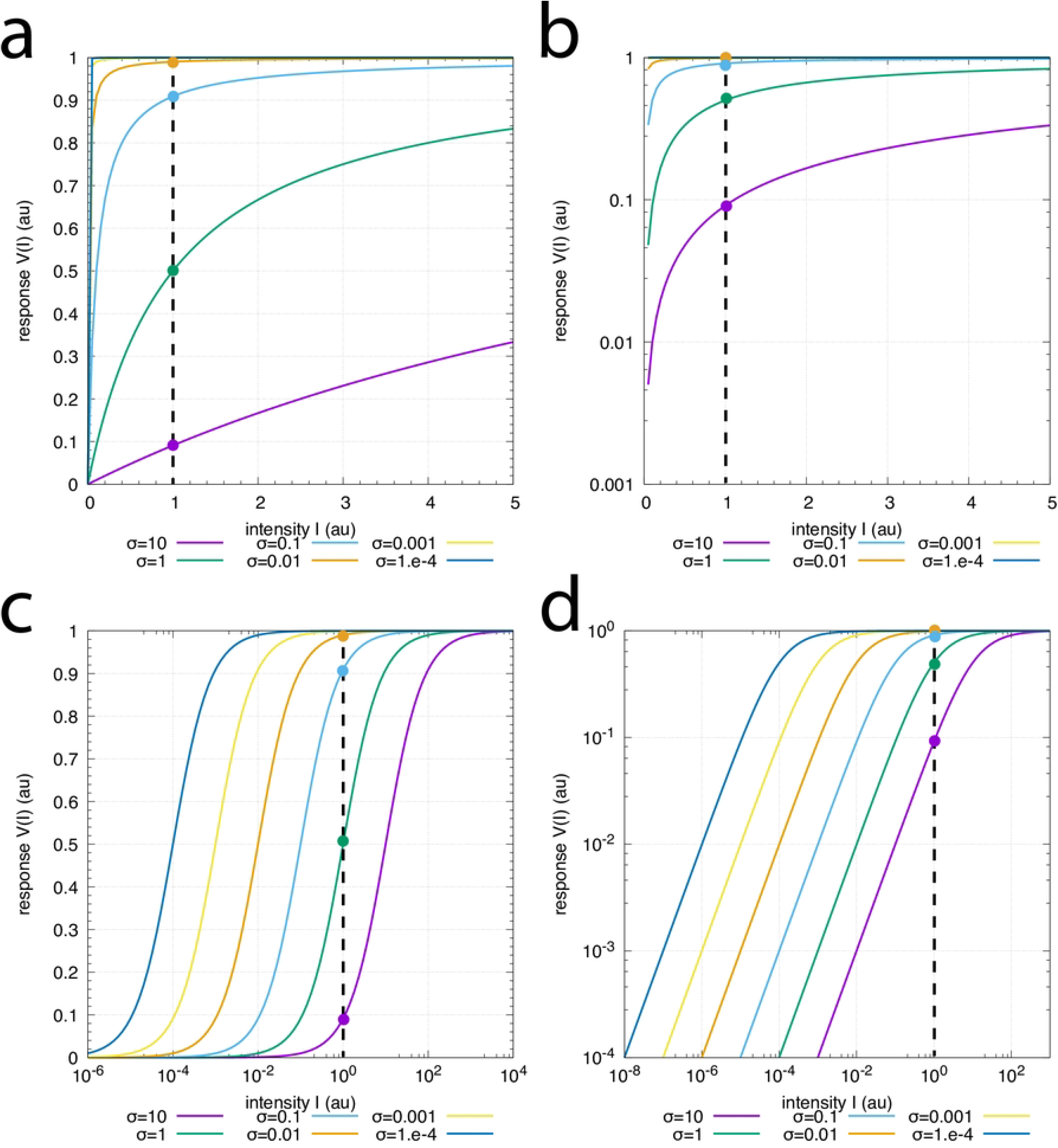
Photoadaptation behaviors in rods and cones described by the Michaelis-Menten equation. The colored lines in the panels show Eq 33 with *σ* values ranging over six orders of magnitudes from *σ*=1*×*10^*−*4^ up to *σ*=10. For simplicity, *V*_*max*_=1. Panel a: Both axes are linear. Panel b: Ordinate is logarithmic and abscissa is linear. Panel c: Ordinate is linear and abscissa is logarithmic. Panel d: Both axes are logarithmic. The dashed vertical lines indicate an perturbation intensity of *I*=1. The colored intersection points with the vertical dashed lines show the responses of *V* for the different backgrounds with the same color.

Fig 18c is analogous to the results in Fig 2 when for a given perturbation (indicated by the vertical dashed lines) an increased background or an increased *σ* leads to a reduction in the averaged maximum frequency. No background compensation, as indicated by Kandel et al. in Ref [2] appears necessary.

When studying the photoadaptation of gecko photoreceptors, Kleinschmidt and Dowling [33] showed log-log relationships analogous to Fig 18d. Dowling interpreted the parallel lines as follows: *A second adaptive mechanism in the receptor shifts the photoreceptor intensity-response curves along the intensity axis, thus extending the range over which the receptor responds* (cited from Ref [3], page 222, bottom section).

Clearly, as Fig 18 shows, the parallel lines in panels c or d neither require the need for a compensation mechanism of a background or other additional adaptive mechanisms. While adaptation mechanisms compensating for a background cannot be excluded, the observation of parallel lines in semi-logarithmic or double-logarithmic plots appear not sufficient to indicate additional background compensation mechanisms besides the negative feedbacks, which lead to the responses in Fig 1 [5].

## Conclusion and outlook

We have shown how robust background compensation in oscillatory and non-oscillatory homeostatic controllers can be realized. The needed feedback condition has been termed ‘coherent feedback’ in analogy to a corresponding concept applied in quantum control theory. Although the property of robust background compensation appears interesting, we are presently not aware of any biological or biochemical example that shows or applies this property. Background compensation may become of interest in synthetic biology to design cellular responses, which by some reason are needed to become background independent. Concerning the case of retinal light adaptation, parallel lines in semi-logarithmic or double-logarithmic plots do not necessarily imply the presence of background compensating mechanisms as defined in this paper.

## Supporting information

### S1 Programs. Documentation

A zip-file with python scripts describing the results for Figs 5, 6, 9a, 9b, 12b, 12c, 14c, 14d, and 16.

